# Connectivity analyses of bioenergetic changes in schizophrenia: Identification of novel treatments

**DOI:** 10.1101/338392

**Authors:** Courtney R. Sullivan, Catharine A. Mielnik, Sinead M. O’Donovan, Adam J. Funk, Eduard Bentea, Erica A.K. DePasquale, Zhexing Wen, Vahram Haroutunian, Pavel Katsel, Amy J. Ramsey, Jarek Meller, Robert E. McCullumsmith

## Abstract

We utilized a cell-level approach to examine glycolytic pathways in the DLPFC of subjects with schizophrenia (n=16) and control (n=16) subjects and found decreased mRNA expression of glycolytic enzymes in pyramidal neurons, but not astrocytes. To replicate these novel bioenergetic findings, we probed independent datasets for bioenergetic targets and found similar abnormalities. Next, we used a novel strategy to build a schizophrenia bioenergetic profile by a tailored application of the Library of Integrated Network-Based Cellular Signatures data portal (iLINCS) and investigated connected cellular pathways, kinases, and transcription factors using Enrichr. Finally, with the goal of identifying drugs capable of “reversing” the bioenergetic schizophrenia signature, we performed a connectivity analysis with iLINCS and identified peroxisome proliferator-activated receptor (PPAR) agonists as promising therapeutic targets. We administered a PPAR agonist to the GluN1 knockdown model of schizophrenia and found it improved long-term memory. Taken together, our findings suggest that tailored bioinformatics approaches, coupled with the LINCS library of transcriptional signatures of chemical and genetic perturbagens may be employed to identify novel treatment strategies for schizophrenia and related diseases.

## INTRODUCTION

Schizophrenia is a devastating illness that affects over 2 million people in the U.S. and displays a wide range of symptoms (1-6). Cognitive impairment is a hallmark feature of the illness, and antipsychotics have poor efficacy in treating cognitive decline (7, 8). The dorsolateral prefrontal cortex (DLPFC) is a brain region heavily implicated in the pathophysiology of schizophrenia, possibly contributing to defects in working memory (9). A variety of abnormalities have been reported in the DLPFC of schizophrenia patients, including changes in gene expression, cell density, receptor binding, and cerebral blood flow (10-12). There is also accumulating evidence of bioenergetic dysfunction in chronic schizophrenia, which has recently been reviewed and highlights a number of abnormalities associated with glucose metabolism, the lactate shuttle, and bioenergetic coupling (13). There is also evidence for genetic linkage between enzymes that control glycolysis in schizophrenia including 6-phosphofructo-2-kinase/fructose-2,6-bisphosphatase 2 (PFKFB2) and hexokinase 3 (HK3), suggesting that genetic risk for this illness includes bioenergetic substrates (14). Interestingly, studies using cell-level techniques have demonstrated complex bioenergetic changes, including decreases in mitochondrial oxidative energy metabolism genes in dentate granule neurons (n=22) and in layer 3 and 5 pyramidal neurons from the DLPFC (n=36 and n=19)(15-17). Decreases in metabolic transcripts included lactate dehydrogenase A (LDHA), nicotinamide adenine dinucleotide dehydrogenases (NADH), and ATP synthases. We have also recently demonstrated decreases in glycolytic enzymes (HK1, and phosphofructokinase muscle type, PFKM) and glucose transporters (GLUT1 and GLUT3) in pyramidal neurons, but not astrocytes, in the DLPFC of schizophrenia subjects (n=16)(18). While examining the expression of individual targets is important, employing a signature based bioinformatics approach may help elucidate the pathophysiology of large biological networks.

Complex illnesses such as schizophrenia are rarely a consequence of an abnormality in the expression of a single gene, but rather a combination of abnormalities in complex cellular networks. This study employs a novel strategy to derive and analyze molecular signatures of schizophrenia by using the Library of Integrated Network-based Cellular Signatures (LINCS). LINCS is an online reference library of cell-based perturbation-response “signatures” of both chemical (small drug-like molecule) and genetic (gene knockdown) perturbations, employing a wide range of assay technologies cataloging diverse cellular responses at the transcriptome, proteome and other levels (19). This reference library used here includes data on the expression of 978 landmark genes (L1000 genes) following the genetic knockdown of genes in various cell lines (with the changes in the L1000 genes from a single genetic perturbation called “knockdown signatures”). The rationale is that the selected landmark genes would capture most of the information contained within the entire transcriptome, allowing researchers to examine disease associated changes at the network level (20). Over 100 cell types have been used to build the LINCS database, and model systems include proliferating immortal cell lines, primary cultures, and induced pluripotent stem cells (iPSCs)(19).

While other bioinformatics resources are also used, our strategy heavily utilizes the iLINCS web portal for querying and complex analyses of LINCS data (http://ilincs.org) to systematically explore the biological underpinning of schizophrenia associated gene expression changes and identify candidate therapeutic strategies from network based molecular changes (Figure 1). We began by modeling a schizophrenia bioenergetic profile in iLINCS based on two new cell-level findings from this study (PFK liver type, PFKL, and glucose-6-phosphate isomerase, GPI), previously identified gene expression changes from cell-level studies (HK1, PFKM, LDHA)(15, 18), and a published genetic linkage study (PFKFB2)(14). The genes comprising our schizophrenia bioenergetic profile are involved in both glycolysis and the lactate shuttle, and are referred to as “seed genes” (Table 1). iLINCS may be queried for each seed gene to determine if a knockdown signature has been catalogued in the database. By clustering the seed gene knockdown signatures in iLINCS, we are able to analyze the connectivity of the knockdown signatures (and thus establish a schizophrenia bioenergetic profile), identifying groups of L1000 genes that change concordantly following genetic perturbation. Furthermore, we can perform Enrichr analyses on these genes to gain perspective on bioenergetic pathology, describe potential multisystem pathological mechanisms, and inform pharmacological studies.

**Figure 1.**
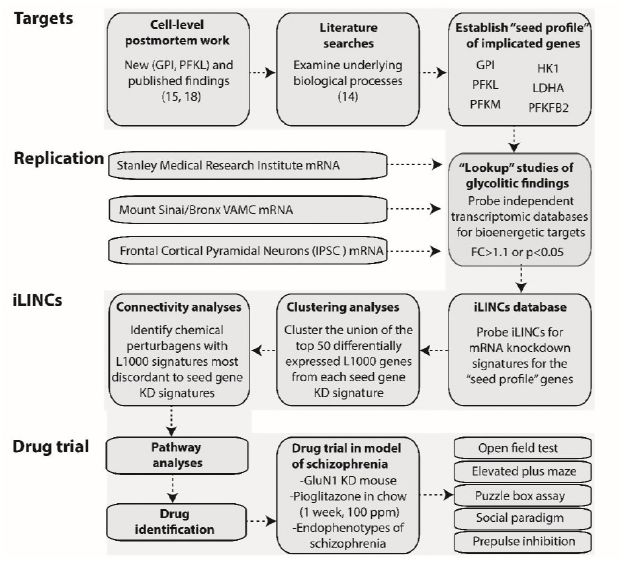
Overview of bioinformatic workflow. We began by performing literature searches and further characterizing cell-level postmortem bioenergetic abnormalities in schizophrenia. Based on these findings, we established a bioenergetic seed profile containing 6 glycolytic genes implicated in schizophrenia ("seed genes”). To replicate our bioenergetic findings in schizophrenia, we probed a publicly available microarray database (Stanley Medical Research Institute Online Genomics Database), microarray data from the DLPFC of control and schizophrenia subjects (Mount Sinai School of Medicine, Bronx VAMC mRNA dataset), and RNAseq data from frontal cortical neurons of a patient with schizophrenia and DISC1 mutation (75). Next, we used iLINCS (http://ilincs.org) to retrieve knockdown signatures for each seed gene individually. With the goal of identifying panels of correlated transcriptomic changes in our seed gene KD signatures, we used iLINCS to perform unsupervised clustering of the top 50 differentially expressed L1000 genes from our 6 seed gene KD signatures. The panels of clustered differentially expressed L1000 genes were carried forward to Enrichr analyses. We also performed iLINCS connectivity analyses and identified peroxisome proliferator-activated receptor (PPAR) agonists as promising therapeutic targets with the potential to “reverse” glycolytic deficits as well as the associated changes in the L1000 genes. Finally, we administered the PPAR agonist pioglitazone to the GluN1 knockdown model of schizophrenia an examined endophenotypes of schizophrenia. Phosphofructokinase liver type (PFKL); PFK muscle type (PFKM); Glucose-6-phosphate isomerase (GPI); 6-phosphofructo-2-kinase (PFKFB2); lactate dehydrogenase A (LDHA); hexokinase 1 (HK1); fold change (FC); inducible pluripotent stem cells (IPSC); knockdown (KD).

**Table 1.**
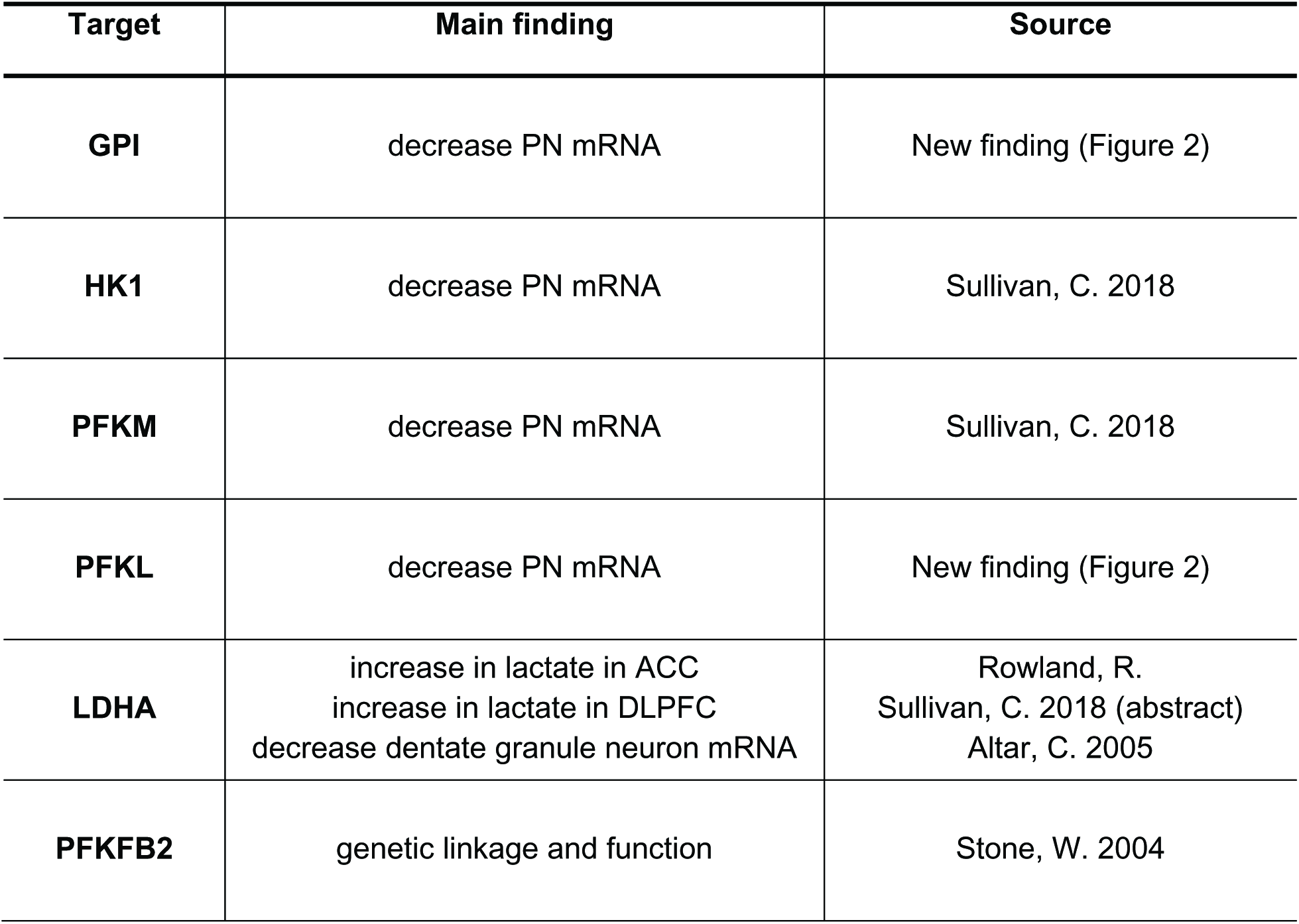
Seed profile for iLINCS analyses. Summary of metabolic findings from cell-level human postmortem studies and literature. Pyramidal neuron (PN), glucose phosphate isomerase (GPI), hexokinase 1 (HK1), phosphofructokinase muscle and liver type (PFKM, PFKL), lactate dehydrogenase A (LDHA), 6-phosphofructo-2-kinase/fructose-2,6-bisphosphatase 2 (PFKFB2).

| Target | Main finding | Source |
| --- | --- | --- |
| **GPI** | decrease PN mRNA | New finding (Figure 2) |
| **HK1** | decrease PN mRNA | Sullivan, C. 2018 |
| **PFKM** | decrease PN mRNA | Sullivan, C. 2018 |
| **PFKL** | decrease PN mRNA | New finding (Figure 2) |
| **LDHA** | increase in lactate in ACC<br>increase in lactate in DLPFC<br>decrease dentate granule neuron mRNA | Rowland, R.<br>Sullivan, C. 2018 (abstract)<br>Altar, C. 2005 |
| **PFKFB2** | genetic linkage and function | Stone, W. 2004 |

Next, we aimed to identify therapeutic compounds with the potential to “reverse” glycolytic deficits as well as the associated changes in the L1000 genes. In order to select a potential drug intervention for preclinical testing, we relied on the ‘connectivity’ analysis to reveal positively (concordant) and negatively correlated (discordant) chemical (small molecule) and genetic perturbations. We developed a discovery based workflow to generate a list of small molecule perturbagens with L1000 transcript changes in the opposite direction of our knockdown signatures (based on concordance values from iLINCS, i.e. generating an “inverse/discordant signature”) (Figure 1). iLINCS provides over 40,000 transcriptomic profiles (signatures) of cell lines following treatment with chemical perturbagens such as FDA approved drugs, chemical probes, and screening library compounds including those with clinical utility and known mechanisms of action (19). FDA approved drugs generated from this “knockdown signature connectivity” analysis could serve as candidates for therapeutic treatment in a preclinical model of schizophrenia. Peroxisome proliferator-activated receptor (PPAR) agonists had multiple hits in our analysis and similar drugs in this class are currently FDA approved for use in humans (21).

One of the leading hypotheses of the etiology of schizophrenia is glutamatergic hypofunction (22-24). N-methyl-D-aspartate subtype glutamate (NMDA) receptor dysfunction is strongly implicated in the pathophysiology of schizophrenia, particularly in human pharmacological studies (25-28). Converging evidence suggests that NMDA receptor hypofunction may actually reflect a dysregulation at the receptor level, including mutations in NMDA receptor subunits (29). Several studies have found associations between the gene that encodes the GluN1 subunit of the NMDA receptor (*Grin1*) and schizophrenia (30-38). The GluN1 knockdown mouse model has a ∼90% global reduction of normal functioning NMDA receptors, achieved by targeted insertion of 2 kb of foreign DNA into intron 19 of the Grin1 gene (39). Additionally, mice with mutations in the Grin1 gene have well-characterized behavioral endophenotypes for schizophrenia; this includes impairments in executive function and working memory, increased stereotypic behavior, decreased anxiety-like behavior, decreased sensorimotor gating, and abnormal social interaction (40). Several of these behaviors are attenuated with typical and atypical antipsychotics at doses similar to those used in pharmacological models of schizophrenia (MK-801 or phencyclidine models), including hyperactivity, stereotypy, and sociability deficits (39, 41). Therefore, the GluN1 knockdown mouse model is a viable and useful tool for studying the interface of NMDA receptor hypofunction, bioenergetics, and schizophrenia symptomology.

There is evidence of abnormal bioenergetic pathways in GluN1 knockdown animals and other models of impaired synapses. For example, there is strong evidence linking lactate shuttle defects to NMDA receptor dysfunction in MK-801 treated rats and in GluN1 subunit knockdown mice (42-44). Dysregulation of the energetic flow of lactate inhibits long term potentiation (45, 46), and may contribute to cognitive deficits in GluN1 knockdown mice. Additionally, when lactate flow from astrocytes to neurons is stimulated, or when lactate is introduced to extracellular space, it enhances working memory and memory formation in rats (45-47). Thus, drug-induced stimulation of lactate production coupled with behavioral studies in GluN1 knockdown mice could help elucidate the role of bioenergetic abnormalities in cognitive dysfunction in schizophrenia.

Pioglitazone is a member of the thiolazinedione (TZD) drug family (that appeared in our knockdown signature connectivity analysis) that is approved to treat type 2 diabetes and hyperglycemia (48). Pioglitazone is a synthetic ligand for PPARγ, a nuclear receptor that is responsible for the regulation of several bioenergetic functions such as lipid homeostasis, adipocyte differentiation, and insulin sensitivity (48, 49). Interestingly, PPARγ agonists have the ability to reduce oxidative stress (via mediating nitric oxide production) as well as modifying mitochondrial metabolism by interacting with proteins associated with mitochondrial function (mitoNEET proteins)(49-53). Activation of PPARγ via pioglitazone can also alter the transcription and expression of GLUT1, leading to changes in glucose uptake through PPARγ and other mechanisms (48, 54). Thus, we hypothesize that treatment with pioglitazone will help restore cognitive endophenotypes associated with schizophrenia in the GluN1 knockdown model via stimulation of metabolic pathways.

In our final experiment, we treated animals with pioglitazone and measured various behavioral outputs that are endophenotypes for schizophrenia (55, 56). First, we assessed locomotor activity, stereotypic behaviors, and mania-like behavior. Next, we assessed executive function and cognitive flexibility using the puzzle-box test. This test requires the mouse to display problem-solving behavior as well as short- and long-term explicit memory to reach a goal area. To assess pioglitazone’s effect on sociability, we used an age- and sex-matched mouse as a social stimulus (57). Finally, we assessed sensorimotor gating using the pre-pulse inhibition (PPI) of acoustic startle paradigm (58). Taken together, these studies provide a bioinformatics translational approach to identify novel treatment strategies through the use of the LINCS library of signatures and connectivity analysis, coupled with the analyses of primary data and published work (to extend the set of ‘seed’ genes).

## RESULTS

### Laser capture microdissection-quantitative polymerase chain reaction studies

We used laser capture microdissection coupled with quantitative polymerase chain reaction (LCM-qPCR) to assess mRNA expression of our metabolic targets in enriched populations of astrocytes and pyramidal neurons in schizophrenia and control subjects (Table S1 and S2). We have previously shown our LCM samples are enriched for specific cell-subtypes using neurochemical markers (11, 18, 59, 60). No significant associations were found between pH, post-mortem interval (PMI), RNA integrity number (RIN), or age and any of our dependent measures. In a sample enriched for pyramidal neurons, we found decreases in PFKL (27%, p=0.010) and GPI (26%, p=0.015) mRNA expression (Figure 2, p<0.05). We did not detect any significant changes in mRNA expression in samples enriched for astrocytes (Figure S1). To assess the use of antipsychotic medication in our patient population, we reanalyzed our significant cell-level targets with and without antipsychotic naïve subjects. All findings remained significant without these subjects (PFKL p=0.037, GPI p=0.034) (Figure S2, panels A and B).

**Figure 2.**
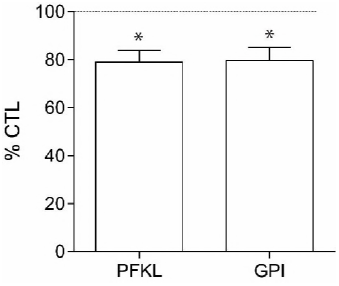
mRNA expression in pyramidal neurons. Relative expression levels of glucose-6-phosphate isomerase (GPI) and phosphofructokinase liver (PFKL) transcripts in enriched pyramidal neuron populations from schizophrenia (SCZ, n=16) and control (CTL, n=16) subjects (A). Data from pyramidal neuron enriched samples were normalized to the geometric mean of three housekeeping genes (B2M, PPIA, ACTB). Data are expressed as percent control ± SEM. *P<0.05.

We also assessed the effects of antipsychotics on PFKL and GPI across multiple brain regions in the Stanley Medical Research Institute genomics database and detected an effect on GPI (p=0.006, fold change=1.09), but not PFKL (p=0.09, fold change=1.06). However, this effect was in the opposite direction of our findings, suggesting the decreased expression in pyramidal neurons in schizophrenia is not due to a medication effect. Additionally, we assessed the effects of ethanol use, smoking, and laterality on our dependent measures in schizophrenia subjects via secondary analyses of schizophrenia subjects on and off ethanol, smokers/nonsmokers, and left/right brain and did not detect any effects (Figure S2, panels C-H).

### Generation of schizophrenia bioenergetic profile (seed genes)

We built a schizophrenia bioenergetic gene profile of 6 glycolytic genes based on data from our cell-level postmortem findings (HK1, PFKM, PFKL, GPI) and the literature (LDHA, PFKFB2). These “seed genes” were used in subsequent “lookup” replication studies, iLINCS clustering analyses (followed by Enrichr analysis), and iLINCS connectivity analyses (drug discovery analysis using “inverse/discordant” signatures). Figure 1 summarizes the bioinformatic work flow for both clustering and connectivity analyses.

### “Lookup” replication studies for seed genes

To replicate our bioenergetic findings, we performed *in silico* validation studies (“lookup” studies) using three independent datasets (Stanley Medical Research Institute (SMRI), Mount Sinai School of Medicine (MSSM), and a frontal cortical neuron RNAseq dataset generated from IPSCs of a patient with schizophrenia and DISC1 mutation (61)). Table 2 summarizes findings from this study and previous studies in our laboratory, as well as the mRNA expression of our selected seed genes (LDHA, HK1, PFKM, PFKL, GPI, PFKFB2) in each dataset.

**Table 2.** Summary of experimental changes and “lookup” replication studies. (all data presented as fold change disease versus control). Summary of region and cell-level seed gene findings from this human postmortem study and online databases. Mount Sinai database compares microarray data from the dorsolateral prefrontal cortex in schizophrenia (n=16) and control (n=15) subjects. Stanley Medical Research Institute (SMRI) Genomics database compares microarray and consortium data from schizophrenia (n=50) and control (n=50) subjects from 5 postmortem brain regions (BA6, BA8/9, BA10, BA46, cerebellum). The cortical neuron mRNA is a publicly available dataset comparing mRNA from inducible pluripotent stem cells from schizophrenia and control patients differentiated in frontal cortical neurons. Stanley Medical Research Institute (SMRI) Online Genomics Database, Mount Sinai School of Medicine (MSSM) dorsolateral prefrontal cortex (DLPFC), fold change (FC), glucose phosphate isomerase (GPI), hexokinase 1 (HK1), phosphofructokinase muscle and liver type (PFKM, PFKL), lactate dehydrogenase A (LDHA), 6-phosphofructo-2-kinase/fructose-2,6-bisphosphatase 2 (PFKFB2). All results are presented as diseased state versus control. Red text indicates finding was significant according to p-value cut off of 0.05 and/or fold change cut off of 1.1. # denotes unavailable p statistics due to low subject number. ## denotes multiple affy probes for the same gene (geomean was used to calculate fold change). *P<0.05 and FC>1.10.

| Target | Human DLPFC mRNA | Pyramidal Neurons mRNA | Cortical Neurons mRNA | SMRI Genomics mRNA | MSSM |
| --- | --- | --- | --- | --- | --- |
| LDHA<sup>15</sup> | -1.15 FC, p=0.359<sup>18</sup> | -1.11 FC, p=0.285<sup>18</sup> | -1.64 FC<sup>#</sup> | -1.11 FC, p=0.022 | -1.06 FC, p=0.320 |
| HK1 | 1.10 FC, p=0.397<sup>18</sup> | -1.24 FC, p=0.003<sup>18</sup> | -1.99 FC<sup>#</sup> | 1.12 FC, p=0.065 | -1.01 FC, p=0.831 |
| PFKM | -1.32 FC, p=0.039<sup>18</sup> | -1.43 FC, p=0.0001<sup>18</sup> | 1.24 FC<sup>#</sup> | 1.05 FC, p=0.225 | 1.03 FC, p=0.694 |
| PFKL | NM | -1.27 FC, p=0.011 | -2.00 FC<sup>#</sup> | -1.00 FC, p=0.920 | -1.03 FC<sup>##</sup> |
| PFKFB2<sup>14</sup> | NM | -1.07 FC, p=0.573<sup>18</sup> | -1.09 FC<sup>#</sup> | 1.00 FC, p=0.975 | -1.31 FC<sup>##</sup> |
| GPI | NM | -1.26 FC, p=0.015 | -1.97 FC<sup>#</sup> | -1.01 FC, p=0.659 | -1.01 FC, p=0.878 |

LDHA is significantly downregulated in both the SMRI (−1.11 FC, p=0.022) and frontal cortical neuron mRNA (−1.64 FC, p=0.018) dataset, while PFKFB2 was downregulated in the MSSM microarray dataset (−1.31 FC). Many seed genes were unchanged in the SMRI and MSSM datasets. However, the SMRI database is an aggregate of 5 brain regions and the MSSM dataset utilized blended samples from the whole DLPFC, while our findings are from cell-level studies. This is supported by 5/6 seed genes being above our fold change threshold in the cell-level frontal cortical neuron dataset.

### Generation of seed gene knockdown signatures

We used iLINCS (http://ilincs.org) to retrieve knockdown signatures for each seed gene individually. Each seed gene knockdown signature (GPI knockdown signature, HXK1 knockdown signature, PFKM knockdown signature, PFKL knockdown signature, LDHA knockdown signature, PFKFB2 knockdown signature) is comprised of the transcriptional changes of 978 landmark genes (L1000 genes) when that seed gene is knocked down (Figure S3). Only signatures designated to be reproducible and self-connected ("gold") by the Broad Institute are available. When multiple shRNAs are available for gene knock-down experiments, consensus gene knock-down signatures, as defined by iLINCS are used. To minimize variability, we compared signatures for seed genes using one consistent cell line (vertebral-cancer of the prostate, VCAP).

### iLINCS clustering analyses

For unsupervised clustering analyses, we used iLINCS to cluster the union of the top 50 differentially expressed L1000 genes from each of the 6 seed gene knockdown signatures using Pearson correlation coefficients (Figures 3 and 4). Due to gene overlap between top 50 differentially expressed L1000 genes in seed gene knockdown signatures, this resulted in the clustering of expression data for 260 L1000 genes. We next used the dendrogram function in iLINCS to select panels of mRNAs with increased expression across clustered seed gene knockdown signatures (“panels of clustered upregulated genes”) and with decreased expression across clustered seed gene knockdown signatures (“panels of clustered downregulated genes”). These clustered gene panels (Table S3) were carried forward to two separate Enrichr analyses.

**Figure 3.**
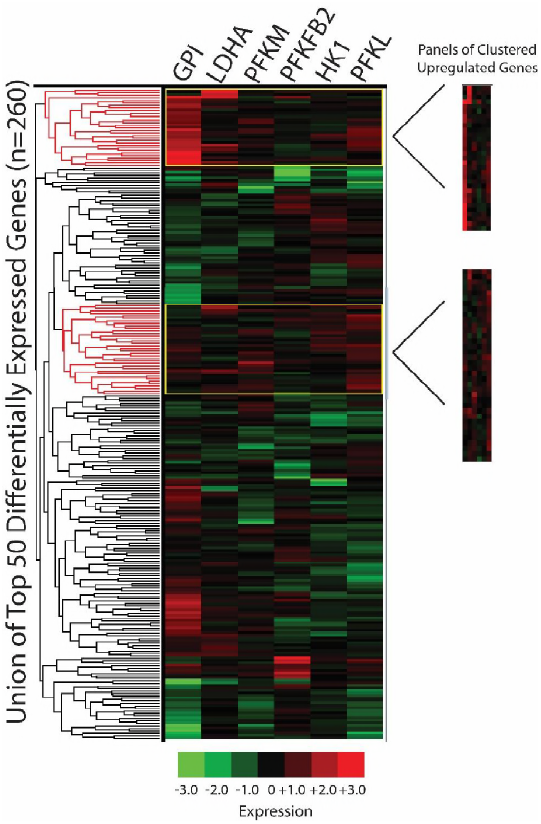
Clustering analyses: panels of clustered upregulated genes. The union of the top 50 differentially expressed L1000 genes per seed gene knockdown signature were clustered in iLINCS via Pearson correlation coefficients (n=260 genes due to gene overlap). Panels of upregulated gene clusters were selected using the dendrogram clustering function and fold change/p-values were exported to Excel. Phosphofructokinase liver type (PFKL); PFK muscle type (PFKM); G lucose-6-phosphate isomerase (GPI); 6-phosphofructo-2-kinase (PFKFB2); lactate dehydrogenase A (LDHA); hexokinase 1 (HK1).

**Figure 4.**
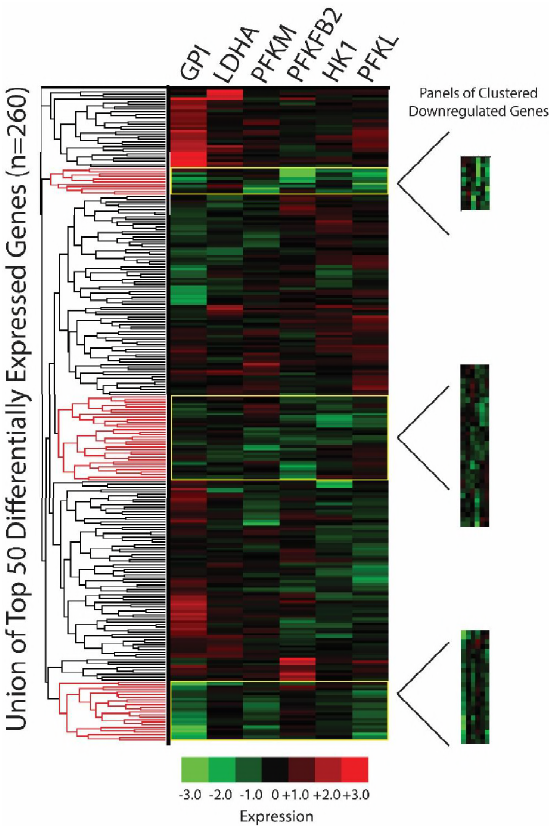
Clustering analyses: panels of clustered downregulated genes. The union of the top 50 differentially expressed L1000 genes per seed gene knockdown signature were clustered in iLINCS via Pearson correlation coefficients (n=260 genes due to gene overlap). Panels of clustered downregulated genes were selected using the dendrogram clustering function (based on Pearson correlation coefficient) and fold change/p-values were exported to Excel. Phosphofructokinase liver type (PFKL); PFK muscle type (PFKM); G lucose-6-phosphate isomerase (GPI); 6-phosphofructo-2-kinase (PFKFB2); lactate dehydrogenase A (LDHA); hexokinase 1 (HK1).

### Enrichr analyses of panels of clustered upregulated genes

We performed Enrichr analyses on panels of clustered upregulated genes from our iLINCS clustering analysis. Top cell signaling pathways included forkhead box O (FOXO), mitogen-activated protein kinase (MAPK), and Wnt signaling pathways (Figure S4). These pathways often interact and all have important roles in cell metabolism, inflammation, cellular proliferation, oxidative stress prevention, and apoptosis (62-65). Analysis of gene ontology (GO) cellular components for our panels of clustered upregulated genes included several microtubule and cytoskeletal components (Figure S5), while the top hits for GO molecular function included protein kinase C (PKC) and MAPK binding (Figure S6). Our protein-protein interactions analysis (Figure S7) yielded multiple hits for protein kinases such as RPS6KA3, MAPK14, GSK3B, and CDK1. We also found hits for proteins involved in protein degradation and protein ubiquitination (CUL1 and SKP1), chromatin remodeling (PCNA and HDAC1), and an estrogen receptor (ESR1).

We next probed for kinases that when knocked down in various cell lines, have upregulated genes similar to the panels of clustered upregulated genes from our iLINCS clustering analysis. Top hits included several serine/threonine-protein kinases (AKT1, MAP2K2, PTK2, ATR) as well as G-protein coupled receptors (GPCRs) involved in energy homeostasis (NMUR2, FFAR1) (Figure S8). We next probed for transcription factors that had occupancy sites for our panels of clustered upregulated genes. We found several transcription factors that play important roles in developmental processes, such as WT1, TRIM28, NFIB, NUCKS1, and TCF3 (Figure S9). Finally, our database of genotypes and phenotypes (dbGaP) analysis implicates diabetes as the most relevant phenotype (Figure S10). A summary of these findings can be found in Table 3.

**Table 3.**
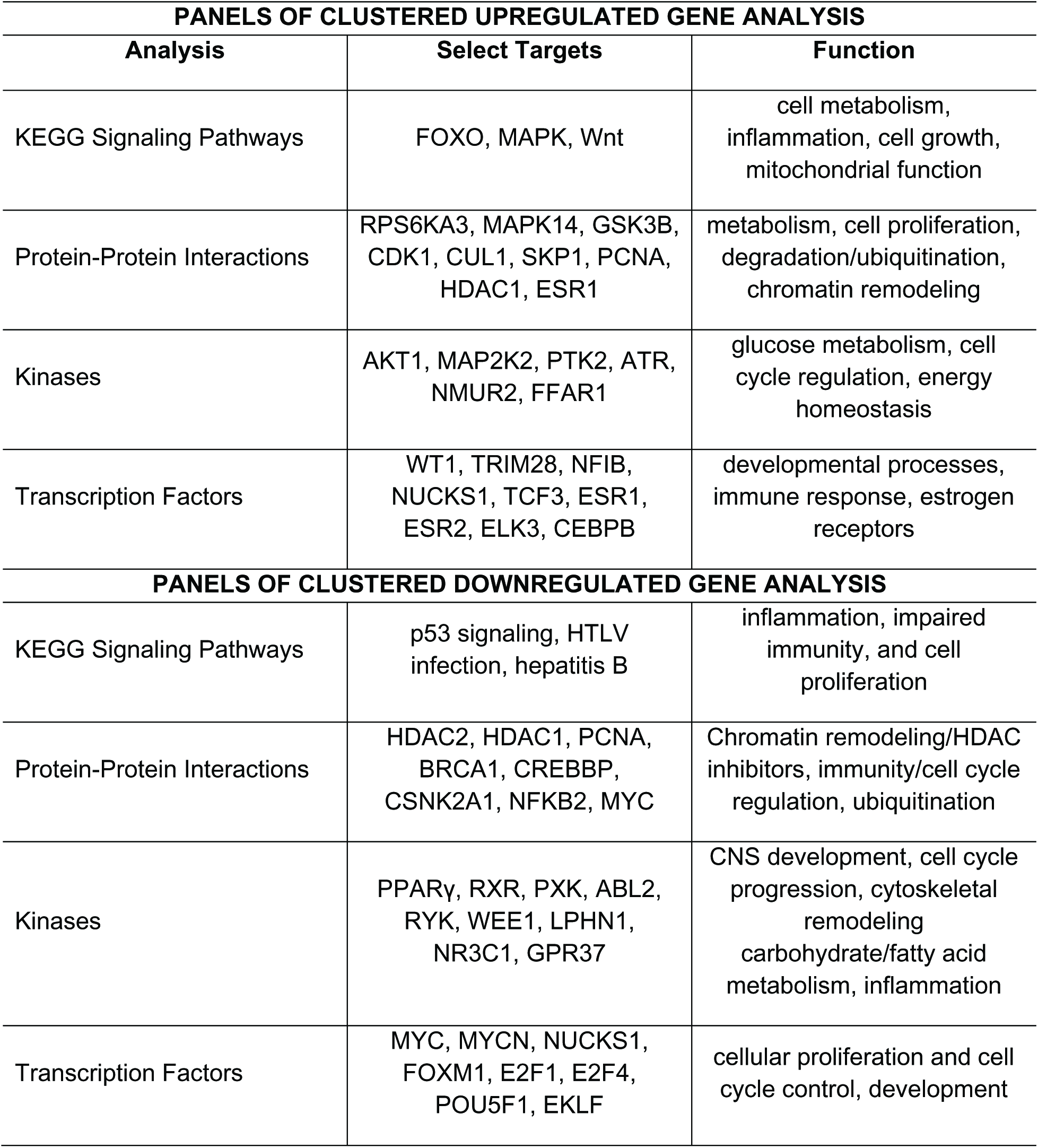
Summary of Enrichr analyses from clustering analysis. Hits and biological function for Enrichr analyses of panels of clustered upregulated and downregulated genes from iLINCS clustering analysis.

### Enrichr analyses of panels of clustered downregulated genes

We performed Enrichr analyses on our panels of clustered downregulated genes from our iLINCS clustering analysis. Top cell signaling pathways include pathways relevant to immune/inflammatory responses and cell cycle regulation (Figure S11). The top 5 hits in our GO cellular components analysis for our panels of clustered downregulated genes implicated mitochondrial function (Figure S12), while the top hits for GO molecular function included several RNA polymerase promotors/activators (Figure S13). Our protein-protein interactions analysis (Figure S14) yielded multiple hits for chromatin remodeling proteins (PCNA, HDAC1, HDAC2, CREBBP). We also found hits for proteins involved in immunity/cell cycle regulation (NFKB2, CSNK2A1, MYC), and ubiquitination (BRCA1).

We next probed for kinases that when knocked down in various cell lines, have downregulated genes similar to our panels of clustered downregulated genes from our iLINCS clustering analysis. Top hits included several kinases involved in fatty acid metabolism and energy homeostasis (RXRA, PPARG, PXK), cellular growth (RYK, WEE1, ABL2), GPCRs (GPR37, LPHN1), and a glucocorticoid receptor (NR3C1) (Figure S15).

We next probed for transcription factors that had occupancy sites for our panels of clustered downregulated genes. We found several transcription factors that play important roles in cell proliferation (MYCN, MYC, E2F1, E2F4, FOXM1), development (NUCKS1, POU5F1), and erythroid function (EKLF) (Figure S16). Finally, our dbGaP analysis implicates carcinoma, memory, and Grave’s Disease as the most relevant phenotypes (Figure S17). A summary of these findings can be found in Table 3.

### iLINCS connectivity analyses (drug discovery)

Using the connected signatures function in iLINCS, we identified chemical perturbagens that produce L1000 signatures that are highly discordant (anti-correlated) with our seed gene knockdown signatures. We generated concordance values (defined in terms of the Pearson correlation coefficients for the gene expression vectors representing the signatures) for every perturbagen in the database, comparing each seed gene knockdown signature to each perturbagen signature separately. The top 20 discordant perturbagen for each seed gene knockdown signature are shown in Table S4 (GPI and PFKM had less than 20 discordant chemical perturbagens, PFKFB2 had none). HK1 and PFKL generated the top 20 perturbagens with the highest discordance (average of −0.4). LDHA, GPI, and PFKM had perturbagen signatures with an average discordance of −0.2 to −0.3. PFKFB2 had no significantly discordant perturbagen signatures.

Next, we combined all discordant perturbagens from Table S4 and reported the top 20 overall discordant perturbagen signatures across all seed gene knockdown signatures (Table S5). After removing duplicate hits, 12 unique perturbagens remained (Table 4). Hits included PPARγ agonists (troglitazone and genistein), valproic acid (a voltage-gated sodium channel blocker and HDAC inhibitor), typical antipsychotic drugs (trifluoperazine, thioridazine, and fluphenazine), and two PI3K inhibitors (Wortmannin and LY-294002). We selected the FDA approved PPARγ agonist pioglitazone for preclinical trials due to its ability to regulate glycolytic pathways that are abnormal in schizophrenia, as well as evidence that pioglitazone treatment improves negative symptoms in patients (cognition has not been studied)(54, 66).

**Table 4.**
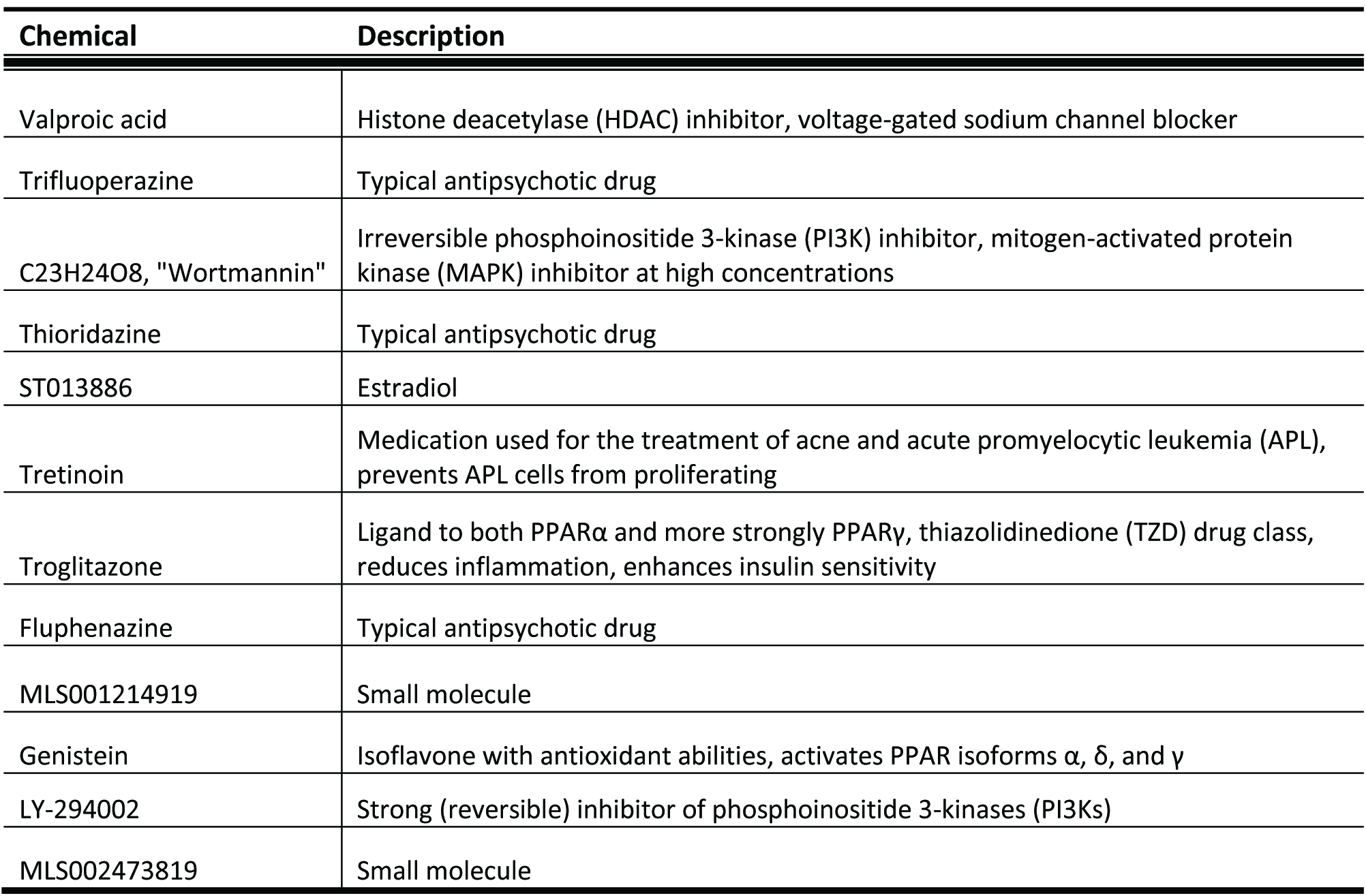
Top 12 unique chemical perturbagens. The top unique chemical perturbagens with the ability to “reverse” the schizophrenia bioenergetic profile.

### Confirming bioenergetic defects in GluN1 knockdown mice

In a cohort of mice separate from our pioglitazone studies, we used a mouse anti-PSD-95 antibody to capture PSD-95 protein complexes from samples (3 male WT, 3 male GluN1 knockdown). The affinity purified samples were enriched for PSD95 when compared to the supernatant and total homogenate (Figure 5A). Following mass spectrometry on affinity purified samples, we analyzed the top 20 increased proteins in GluN1 knockdown mice relative to WT mice with Ingenuity Pathway Analysis (IPA). The top 5 implicated pathways in the GluN1 knockdown mice were gluconeogenesis, glucose metabolism, pyruvate and citric acid metabolism, metabolism of carbohydrates, and glycogen storage disease (Figure 5B). Finally, we examined mRNA expression of metabolic transporters (MCT1, MCT2, MCT4, GLUT1, GLUT3) using real time quantitative polymerase chain reaction (RT-qPCR) in a cohort of GluN1 knockdown and wildtype (WT) mice (n=5 per group). We found significant decreases in GLUT1 (47%, p=0.018) and GLUT3 (34%, p=0.013) in GluN1 knockdown mice compared to WT controls (Figure 5C), confirming that a strategy to increase glucose uptake capacity might be a successful intervention for schizophrenia.

**Figure 5.**
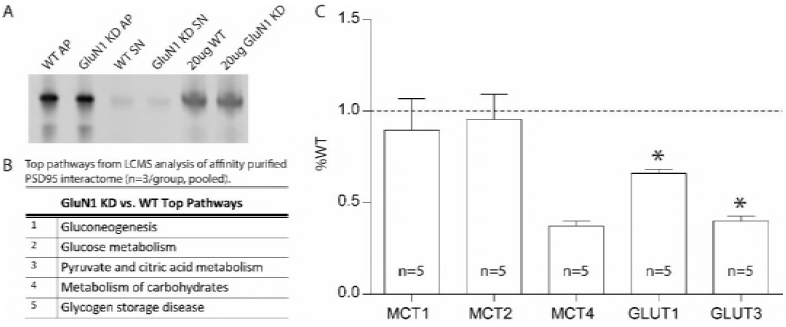
Metabolic defects in the GluN1 knockdown mouse model. Representative PSD-95 Affinity Purification of WT and GluN1 KD mouse samples (A). Enrichr pathway analysis of the top 20 interacting proteins for postsynaptic density 95 (PSD95) interactome for GluN1 KD versus WT mice (n=3/group, pooled) (B). Metabolic transporter transcripts in frontal cortex of GluN1 KD mice expressed as percent wildtype (WT) (n=5) (C). Wildtype (WT); affinity purification (AP); knockdown (KD); supernatant (SN); liquid chromatography-tandem mass spectrometry (LCMS); monocarboxylate transporter (MCT); Glucose transporter (GLUT); Wildtype (WT). Data are expressed as percent control ± SEM. *p<0.05.

### GluN1 knockdown pioglitazone studies

For pioglitazone studies (in a different cohort of mice, Table S6), we assessed caloric intake and body weight to ensure pioglitazone treatment did not have adverse effects. Pioglitazone did not affect the intake of chow or the body weight of the animals (p=0.5096) (Figure S18).

In the puzzle box assay, we found that one week of pioglitazone treatment decreased (improved) time to perform task in GluN1 mice in Trial 1 (training, p=0.011) and Trial 4 (long-term memory, p=0.017) (Figure 6, Panels C and D).

**Figure 6.**
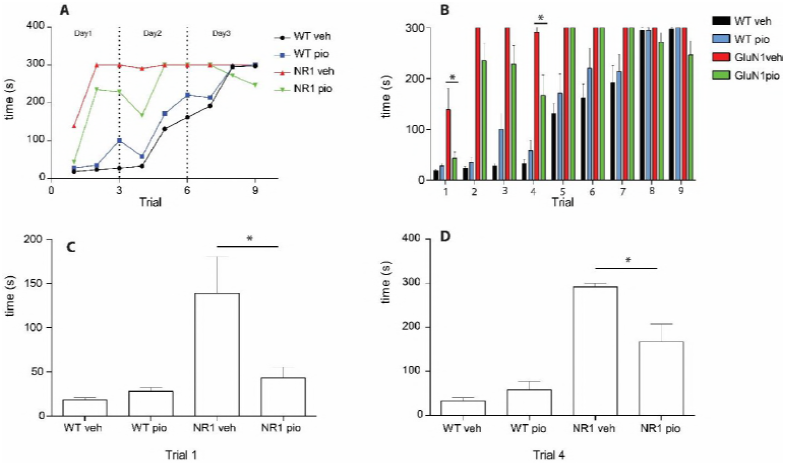
Puzzle box assay. Latency (s) to complete trials in puzzle box expressed as a time course (significance not denoted) (A). Latency (s) to goal zone in puzzle box in WTveh (n=12), WTpio (n=10), GluN1veh (n=9), and GluN1pio (n=12) mice (B). Trial 1 of the puzzle box assay demonstrating a decrease in latency to goal zone in GluN1pio mice when compared to GluN1veh (C). Trial 4 of the puzzle box assay demonstrating a decrease in latency to goal zone in GluN1pio mice when compared to GluN1veh (D). Only drug significance (not genotype) is denoted. Data are shown as mean ± SEM. p<0.05.

In the open field test, we detected a genotype effect on locomotor activity, stereotypy, and vertical activity (p< 0.0001, F (1, 23) = 107.3, F (1, 23) = 363.4, and F (1, 23) = 34.00 respectively) (Figure 7, Panels B-D). GluN1 knockdown mice, when compared to WT littermates, displayed an increase in hyperlocomotion, as quantified by an increase in total distance traveled measured in centimeters, which was accompanied by an increase in overall stereotypic movements and rearing. We did not detect changes in hyperactivity, stereotype, or vertical activity in GluN1 knockdown mice following pioglitazone treatment (Figure 7, Panels BD).

**Figure 7.**
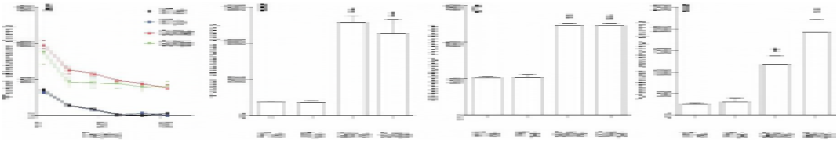
Locomotor activity and stereotypy. Total distance (cm) over time (mins) in the open field test (significance not denoted) (A). Total distance (cm) traveled during open field test (B). Stereotypy number for each group in open field test (C). * denotes significantly different from WTveh. # denotes significantly different from both WTveh and WTpio groups. Data shown as mean ± SEM. P<0.05.

In the elevated plus maze, we detected an effect of genotype (p< 0.0001). GluN1 knockdown animals spent significantly more time in the open arms than did WT mice, regardless of pioglitazone/vehicle treatment (Figure S19). Increased time in the open arm indicates a lack of anxiety and is observed in mouse models of mania-like behavior. We did not detect any significant differences in genotype or drug treatment in our social paradigm (Figure S20).

Finally, in measures of sensorimotor gating, we detected an effect of genotype and genotype by drug interaction at 4 decibels (dB) (p=0.0173 and p=0.0002). Pioglitazone-treated GluN1 knockdown (GluN1pio) animals had decreased PPI compared to all other groups (Figure 8, Panel A). At 8 dB we see an effect of genotype (p=0.0006). GluN1pio mice have significantly lower PPI than either WT group (Figure 8, Panel A). At 16 dB we also see an effect of genotype (p=0.006). GluN1pio mice have significantly lower PPI than do WTpio mice (Figure 8, Panel A). We did not detect any differences in the acoustic startle response between any groups, although GluN1 knockdown animals tended to have higher startle responses (Figure 8, Panel B).

**Figure 8.**
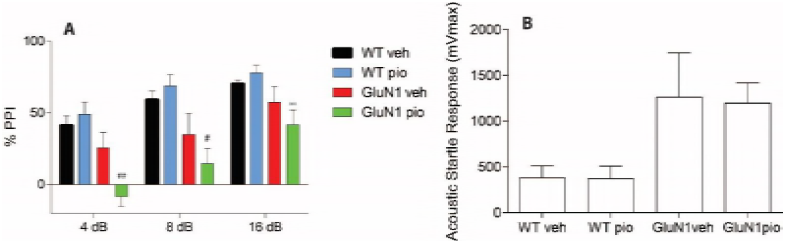
Prepulse inhibition and acoustic startle response. Sensorimotor gating (prepulse inhibition) represented as percent inhibition of acoustic startle response at three prepulse tones; 4db, 8db, and 16dB (A). Acoustic startle response to 165dB startle pulse in pre-pulse inhibition test (B). * denotes significantly different from WTveh; ** denotes significantly different from WTpio; # denotes significantly different from WTveh and WTpio; ## denotes significantly different from WTveh, WTpio, and GluN1veh. Data shown as mean ± SEM. p<0.05.

## DISCUSSION

Big biomedical data resources, such as LINCS, open new avenues for innovative analyses of primary data generated in conjunction with focused, hypothesis driven research being undertaken in individual research labs. Interdisciplinary studies that combine such resources and research expertise in neuroscience, bioinformatics, and data science can greatly enhance our ability to generate novel hypotheses and offer key insights into the mechanisms of human disease. The goal of this study was 4 parts: first, to further characterize changes in glycolytic enzyme expression at the cellular level in schizophrenia and to explore these findings in available *in silico* databases; second, to analyze the connectivity of bioenergetic pathology in schizophrenia by generating a schizophrenia bioenergetic profile in iLINCS and performing pathway analyses on panels of clustered genes that are highly differentially expressed across knockdown signatures; third, to generate a list of promising drug interventions with the goal of future preclinical testing; and fourth, to examine endophenotypes of schizophrenia following treatment with an FDA approved drug identified by our bioinformatic analyses in an animal model.

### Cell-level postmortem studies

Our findings of decreased PFKL and GPI mRNA in pyramidal neurons build upon previous reports of abnormalities in glycolytic enzymes, as well as glucose/lactate transporters, in the DLPFC in schizophrenia (18). PFKL codes for the liver (L) subunit of the PFK enzyme, which catalyzes the phosphorylation of fructose 6-phosphate to fructose 1,6-bisphosphate in the third, irreversible and rate-limiting step of glycolysis (67). There are several tetrameric isoforms of the PFK enzyme, each comprised of different combinations of L, muscle (M), or platelet (P) subunits. The PFK-5 protein is entirely comprised of L subunits, while the PFK-1 protein is entirely comprised of M subunits. Other erythrocyte PFK enzymes contain a heterogenic composition, and all PFK enzymes are found in brain (67). Under conditions of high glycolytic flux, cells can divert glucose to the pentose phosphate pathway; however, the conversion of fructose 6-phosphate to fructose 1,6-bisphosphate by PFK commits glucose to the glycolytic pathway. As a gatekeeper in the metabolic degradation of glucose, PFK is highly regulated, both by downstream metabolites and its strong allosteric activator fructose 2,6-bisphosphate (F2,6BP), which is produced by PFKFB when glycolysis is upregulated (Warburg effect) (68). Notably, knockdown of PFKL impairs glycolysis and promotes metabolism via the pentose phosphate pathway (69). Decreases in PFKL or PFKM expression in schizophrenia could affect normal enzyme activity, resulting in either decreased glycolytic breakdown of glucose or an inability to upregulate glycolysis when energy demand is high. Interestingly, the human PFKL gene is located on chromosome 21 and abnormal expression of PFKL is associated with Down’s syndrome (70). Additionally, we previously reported decreased PFK activity in the DLPFC of schizophrenia subjects (18).

We also found a decrease in the mRNA expression of GPI in an enriched population of pyramidal neurons in schizophrenia. GPI is a dimeric cytoplasmic enzyme that catalyzes the reversible isomerization of glucose-6-phosphate to fructose-6-phosphate in the second step in the glycolytic pathway. When expressed extracellularly, the GPI protein functions as a neurotrophic factor that promotes neuronal survival (71). One study found an increase in glucose-6-phosphate levels, as well as several other metabolites in the glycolytic pathway, in peripheral blood mononuclear cells of schizophrenia patients, suggesting improper functioning of glycolytic enzymes (72). Other data suggest that GPI deficiency results in an accumulation of glucose-6-phosphate and over activation of the mammalian target of rapamycin (mTOR) pathway (73). A decrease in GPI expression in pyramidal neurons could contribute to dysregulation of both glycolysis and mTOR signaling. A disruption in mTOR signaling could affect protein synthesis and lead to aberrant neuronal growth and synapse connectivity. Supporting this hypothesis, the mTOR pathway has been previously implicated in schizophrenia (74).

### Schizophrenia bioenergetic profile (seed genes)

Based on strong cell-level evidence from this study and prior work, as well as decreases in glycolytic enzyme activity, we generated a schizophrenia bioenergetic profile consisting of 6 functionally related genes (14, 15, 18). These 6 genes (“seed genes”) are strong glycolysis regulators and critical steps in the glycolytic pathway. The 6 seed genes in our profile were used for lookup replication studies, iLINCS clustering, and iLINCS connectivity (drug discovery) analyses.

### “Lookup” replication analyses

We sought to replicate our metabolic LCM findings, as well as explore the mRNA expression of seed genes through *in silico* “lookup” analyses (18). In our LCM studies, we found decreases in HK1, PFKM, PFKL, and GPI in pyramidal neurons in the DLPFC in schizophrenia (Figure 2, Table 2)(18). We probed a second cell-level dataset (frontal cortical neurons differentiated from IPSCs of schizophrenia subjects with the DISC mutation (75)) and found that 5/6 of our seed genes were abnormally expressed (LDHA, HK1, PFKM, PFKL, and GPI) (Table 2)(75). LDHA, which was previously found to be downregulated in dentate granule neurons in schizophrenia, was also significantly downregulated in the SMRI genomics database (15). We also found that PFKFB2, which is genetically linked to schizophrenia, is downregulated in schizophrenia in the MSSM DLPFC microarray data (14). The SMRI and the MSSM databases contain region-level findings in multiple brain regions, while IPSCs and LCM techniques provide cell-subtype specific resolution. Further, prior work in our laboratory found a decrease in PFK and HK enzyme activity levels in the DLPFC in schizophrenia (18). These findings suggest our seed genes are dysregulated at the cell, region, transcript, and functional level in schizophrenia, as well as by genetic association.

### Generation of seed gene knockdown signatures

We used iLINCS (http://ilincs.org) to retrieve knockdown signatures for each seed gene individually. Each seed gene knockdown signature (GPI knockdown signature, HK1 knockdown signature, PFKM knockdown signature, PFKL knockdown signature, LDHA knockdown signature, PFKFB2 knockdown signature) is comprised of the transcriptional changes of 978 landmark genes (L1000 genes) when that seed gene is knocked down (Figure S3).

### Panels of clustered upregulated and downregulated gene identification

After clustering the knockdown signatures of our 6 seed genes in iLINCS, we (separately) selected panels of clustered upregulated and downregulated genes using the interactive dendrogram function. This resulted in 67 upregulated and 69 downregulated genes carried forward to Enrichr analyses.

### Enrichr analyses of upregulated genes from iLINCS clustering

We began by using Enrichr to identify pathways associated with our panels of clustered upregulated genes from our iLINCS clustering analysis (Figure 3, Figures S4-S10). Using KEGG, we generated top cell signaling pathways which returned hits such as FOXO, Wnt, and MAPK signaling. These signaling pathways implicate glucose utilization, energy homeostasis, mitochondrial function, and cell growth. Our top protein-protein interaction hits (RPS6KA3, MAPK14, glycogen synthase kinase 3 beta (GSK3B) and cyclin dependent kinase 1 (CDK1), etc.) suggest a role for pathways involved in energy metabolism and cell proliferation. Next, we probed for kinases that when knocked down in specific cell lines, increased the expression of genes that were in our panels of clustered upregulated genes. We found several serine/threonine-protein kinases including AKT1, PTK2, MAP2K2, and ATR. AKT1 is one of 3 closely related serine/threonine-protein kinases (AKT1, AKT2 and AKT3) thought to regulate metabolism and promote cellular proliferation (76). We also had significant hits for G-protein coupled receptor proteins NMUR2 and FFAR1. Neuromedin U (NMUR2) encodes a G-protein coupled receptor protein widely expressed in the central nervous system (CNS) that binds the neuropeptide neuromedin U in order to regulate energy homeostasis (77, 78). These results suggest that kinases involved in glucose metabolism and cell cycle regulation are relevant to our panels of clustered upregulated genes. Finally, we performed pathway analyses to identify transcription factors with occupancy sites for our panels of clustered upregulated genes, several of which played key roles in neurodevelopment (WT1, NFIB, TCF3, TRIM28, etc.). Our transcription factor analysis also generated hits for estrogen receptors (ESR1, ESR2), tumor suppressors/activators (ELK3), and transcription factors involved in immune responses and macrophage function (CEBPB)(79, 80). Enrichr analyses are summarized in Table 3 and a full discussion of panels of clustered upregulated genes can be found in the supplement.

### Enrichr analyses of downregulated genes from iLINCS clustering analysis

We used Enrichr to identify pathways associated with our panels of clustered downregulated genes from our iLINCS clustering analysis (Figure 4, Figures S11-S17). Using KEGG, we generated top cell signaling pathways which returned hits such as HTLV infection, hepatitis B, p53 signaling, and cell cycle pathways (Figure S11). This implicates inflammation, impaired immunity, and cell proliferation. The top 5 hits in our GO cellular components analysis for panels of clustered downregulated genes implicate mitochondria and oxidative phosphorylation (Figure S12). The top hits of our protein-protein interaction analysis (Figure S14) for panels of clustered downregulated genes included histone deacetylase 1 (HDAC1) and HDAC2, a ubiquitously expressed protein that removes acetyl groups from lysine residues on core histones (81). Other top hits in our protein-protein interaction analysis include proliferating cell nuclear antigen (PCNA) and BRCA1, which also bind HDAC1 and are involved in epigenetics, DNA replication, and DNA repair (82). Interestingly, HDAC1 is increased in the prefrontal cortex and hippocampus in schizophrenia and overexpression of HDAC1 leads to impairments in working memory (83-85). There are accumulating instances of schizophrenia patients with mutations in genes encoding chromatin regulators (such as histone modifying enzymes and transcription factors)(86). Other hits included transcription factors involved in inflammation and the balance of activation and suppression of cellular proliferation processes. Next, we probed for kinases that when knocked down in specific cell lines, decreased the expression of genes that were in our panels of clustered downregulated genes (Figure S15). We found multiple kinases with important roles in glucose homeostasis and energy metabolism. The nuclear receptor PPARγ forms a heterodimer with the retinoid X receptor (RXR), two of our top hits, increasing the transcription of various genes that stimulate glucose uptake and carbohydrate metabolism (87). PPARγ ligands such as thiazolidinediones (TZDs) increase glucose utilization, treat hyperglycemia, and represent therapeutic possibilities in diseases with deficient glycolytic systems (88). We also found kinases that are involved in axonal growth during CNS development (RYK), cell growth and cycle progression (WEE1, ABL2), and cytoskeletal remodeling (ABL2). Finally, we performed pathway analyses to identify transcription factors with occupancy sites for our panels of clustered downregulated genes, several of which played key roles in cellular proliferation/development and were similar to hits from previous analyses (MYC, MYCN, NUCKS1, forkhead box M1) (Figure S16). Enrichr analyses are summarized in Table 3 and a full discussion of panels of clustered downregulated genes can be found in the supplement.

In summary, we extrapolated on the abnormalities of glycolytic enzymes using Enrichr pathway analyses for panels of clustered up and downregulated genes. These analyses yield additional pathways and regulators involved in cellular processes such as cell cycle regulation and inflammatory responses. Inappropriate cell cycle regulation could contribute to metabolic deficits, while the immune system has previously been implicated in schizophrenia (89-91). Several of our metabolic and immune findings were replicated in other large scale transcriptomic bioinformatic analyses of psychiatric disorders (92). These results highlight the connectivity of glycolytic pathology in schizophrenia to several systems that could be important targets in future studies.

### iLINCS connectivity analyses (drug discovery)

We divided the top 12 unique perturbagens from our signature connectivity analysis into 5 groups: PPAR agonists, PI3K inhibitors, antipsychotic drugs, HDAC inhibitors, and other (Table 4). PI3K inhibitors constituted 2 of the drugs that may “reverse” our disease signature.

These antitumor drugs inhibit PI3K enzymes, which are part of the PI3K/AKT/mTOR signaling pathway and play critical roles in many cellular functions such as cellular proliferation, metabolism, and immune cell activation (93-95). Recent work has linked schizophrenia pathology to the PI3K-AKT-mTOR signaling cascade (74, 96, 97). Additionally, PI3K signaling can modulate synaptic formation and plasticity, and has been implicated in both schizophrenia and autism disorder (reviewed in (98, 99)). Previous work demonstrated pharmacological inhibition of the catalytic subunit of PI3K blocks amphetamine induced psychosis in mice, highlighting PI3K inhibitors as therapeutic targets for the treatment of psychiatric disorders (100). However, PI3K inhibition may be more suitable for “positive symptoms,” as PI3K inhibition results in diminished insulin-stimulated glucose uptake by inhibiting translocation of GLUT4 glucose transporters to the plasma membrane (101). Three of our drug discovery analysis hits were typical antipsychotics, which also treat positive symptoms of schizophrenia via dopamine receptor D2 antagonism (102). It is possible that the glycolytic knockdown signatures used to generate these perturbagens could cause transcriptional changes in L1000 genes involved in the regulation of dopaminergic synapses, suggesting dopamine and metabolic systems could be connected in schizophrenia. This is not entirely surprising as antipsychotics have well documented metabolic effects (103).

The top unique perturbagen in our signature connectivity analysis (Table 4) was valproic acid, an HDAC inhibitor that modulates sodium channels and enhances gamma-aminobutyric acid (GABA)-mediated neurotransmission (traditionally used to treat epilepsy and bipolar disorder)(104). Interestingly, sodium valproate (valproic acid) is commonly used as an adjunctive therapy for the treatment of schizophrenia, and a recent 4 week randomized clinical trial demonstrated improvement in psychopathology with a combination therapy of valproate and risperidone or olanzapine compared to antipsychotics alone (n=249)(105). There is also some evidence for positive effects on aggression and tardive dyskinesia in schizophrenia, although sample sizes are small (106). Valproic acid also affects ERK and Wnt pathways, which were implicated in our pathway analyses and regulate cell survival and cytoskeletal modifications (107). Stimulation of ERK1/2 has the ability to enhance the transcriptional activity of PPARs and increase glucose metabolism, indicating PPAR agonists might also be potential therapeutic targets (108). Interestingly, two of our unique perturbagen hits included PPAR agonists (troglitazone and Genistein) (109). Troglitazone, part of the TZD family, is an anti-inflammatory and antidiabetic drug developed to treat type 2 diabetes. Troglitazone is a ligand to both PPARα and (more strongly) PPARγ, promotes glucose uptake by increasing two transporters we previously found decreased in pyramidal neurons in schizophrenia (GLUT1 and GLUT3), and inhibits the pro-inflammatory factor NFKB (implicated in our Enrichr analyses)(18, 54, 110, 111). The TZD family of PPAR agonists present an interesting therapeutic intervention for studies in preclinical models of schizophrenia due to their ability to modulate glucose systems we find abnormal in schizophrenia (112, 113). Pioglitazone, a current FDA approved member of the TZD family, also stimulates glycolytic systems and can attenuate mitochondrial dysfunction, reverse memory impairment, and decrease the incidence of dementia (48, 113-119). Thus, pioglitazone is an easily accessible drug candidate to “reverse” pathology and possibly restore cognitive defects related to schizophrenia.

To test this hypothesis, we first examined metabolic pathways in the GluN1 knockdown model of schizophrenia. GluN1 knockdown animals display a wide variety of endophenotypes associated with schizophrenia, including deficits in cognition (40). We found that top pathways associated with protein changes at the synapse in GluN1 knockdown mice compared to WT controls were metabolic in nature (Figure 5B). Additionally, expression of glucose transporters (GLUT1 and GLUT3) were significantly decreased in the frontal cortex of GluN1 knockdown mice (Figure 5C). These results suggest there are similar abnormalities in bioenergetic systems in this model to schizophrenia, and that modulating these systems are viable treatment strategies. Thus, we examined the effects of pioglitazone, a PPAR agonist that increases glucose uptake, on behavioral endpoints in the GluN1 knockdown model. Pharmacological manipulation of glycolytic pathways via pioglitazone could stimulate glucose uptake and metabolism, possibly restoring some behavioral deficits. We are also able to control for medication treatments, which can be difficult to interpret in human studies. It is important to distinguish significant metabolic changes as underlying features of the illness and not epiphenomena related to treatment with antipsychotics.

We hypothesized that pioglitazone treatment would improve cognitive function in the GluN1 knockdown model. The puzzle box assay examines cognitive function and is progressively more difficult, evident in the similarly poor performance by all groups in later trials. However, in all earlier trials (where GluN1 deficit is more robust), pioglitazone tended to increase WT mean latencies while decreasing GluN1 knockdown mean latencies. Our statistical analyses found that in Trial 1 and Trial 4, pioglitazone significantly decreased the latency to perform the task in GluN1 knockdown mice (Figure 6, panels C and D). This suggests that in normal physiology, pioglitazone could have a negative effect on learning, while in a pathological brain with impaired synapses, pioglitazone may have restorative effects on cognition.

We also examined the effects of pioglitazone on behaviors reflecting positive and negative schizophrenia symptoms. In line with published findings for this model (39), our data suggest that GluN1 knockdown animals have increased locomotor activity and stereotypy compared to WT littermate controls, as well as increased mania and anti-anxiety like behavior (increased % time spent in open arms in EPM compared to WT mice) (Figures 7 and S19). Pioglitazone did not have an effect on these behaviors regardless of genotype, suggesting that either glucose metabolism does not play a role in this phenotype of GluN1 knockdown mice or that this is a neurodevelopmental phenotype that requires earlier intervention.

We did not detect any changes in sociability between any groups. All groups choose to spend more time in the social zone (Figure S20). Previous studies demonstrate inherent social abnormalities in GluN1 knockdown mice, suggesting our experiment may be underpowered to detect an effect of sociability (39). We also did not detect any changes in social novelty-however our social novelty data has high variability, which could contribute to lack of significant findings.

Finally, we assessed sensorimotor gating using pre-pulse inhibition and acoustic startle response (Figure 8). While there were no changes in acoustic startle response across all groups, we detected an effect of genotype on PPI at 4 dB (F (1, 23) = 20.06, p=0.0002), 8 dB (F (1, 23) = 15.63, p=0.0006), and 16 dB (F (1, 23) = 9.177, p=0.0060). GluN1 knockdown mice had generally lower PPI than did WT controls at all dBs. Additionally, at 4 dB, pioglitazone-treated GluN1 knockdown (GluN1pio) mice had a lower PPI than any other group, suggesting pioglitazone interacts in a negative way with the knockdown pathophysiology (drug by genotype interaction, F (1, 23) = 6.579, p=0.0173). The 4dB pre-pulse is the lowest level of pre-pulse and thus the “most sensitive” measure of the deficit (vs. 8dB and 16dB, where the deficit is sometimes not seen in a disease state). This could explain why we only detect this negative drug interaction at the 4 dB pre-pulse. Interestingly (although not significant), WTpio mice tended to have a larger PPI versus WTveh mice in all dB tests, while GluN1pio mice tended to have a lower PPI versus GluN1veh mice in all dB tests, reinforcing the idea of different genotype by drug interactions. Additional test animals are needed to explore this possibility.

Our findings suggest that pioglitazone may selectively exert its effects on specific endophenotypes of schizophrenia, suggesting circuit-specific modulation of neuroplastic substrates. One possibility is that pioglitazone improves the function of brain regions and circuits that are hypometabolic, while having no effect/negative consequences on brain regions that are normal or hypermetabolic. Many behavioral endpoints that are markers of “positive symptoms” of schizophrenia could correspond to “hypermetabolic” circuitry, and thus not be amenable to rescue via pioglitazone. We found pioglitazone had no effect on behavioral deficits such as hyperactivity, increased stereotypy, and social impairments, which are normalized by typical and atypical antipsychotics in GluN1 knockdown animals (39, 41, 120-123). However, pioglitazone did have a restorative effect on cognition (puzzle box assay), while typical antipsychotics do not (7, 124). Studies examining the effects of antipsychotic treatment on cognition in GluN1 knockdown animals has not yet been done and are warranted.

Cognition includes several subprocesses, not all of which are uniquely sustained by the frontal cortex, but also by distributed cortical networks including frontal regions which may not be associated with the frontal lobes (125, 126). These processes can include task management, planning, flexibility of behavior and thought, working memory, attention, or others. Thus, poor performance in any of these parameters could reflect a number of possible cognitive mechanisms (125, 127). For instance, two meta-analyses of cannabis use on cognition in schizophrenia demonstrated cannabis improved certain subtasks for elements of cognition (visual memory, planning, working memory) while having no effect or worsening other tasks (attention, verbal memory, processing speed)(128, 129). It is also possible that pioglitazone may only improve a metabolic subtype of schizophrenia. This is supported by our LCMS and qPCR studies in GluN1 knockdown mice showing a metabolic profile. Cognition and other related functions disrupted in schizophrenia are often highly complex, and modulation of these circuits might be treated with personalized combination therapeutics.

For example, one study examined the effects of pioglitazone as an adjunct treatment to antipsychotic drugs in schizophrenia subjects (n=40)(66). Subjects received risperidone plus either pioglitazone (30 mg/day) or placebo for 8 weeks. Patients in the pioglitazone group showed significantly more improvement in Positive and Negative Syndrome Scale (PANSS) negative subscale scores (p < 0.001) as well as PANSS total scores (p = 0.01) compared with the placebo group, suggesting pioglitazone and risperidone work through different circuitry and could be used in conjunction to treat both cognitive/negative and positive symptoms. There is also evidence suggesting pioglitazone treatment could help combat glucose–lipid metabolic abnormalities and diabetes that are often exacerbated by antipsychotics (119). These findings suggest the possibility of pioglitazone as an augmentation therapy in reducing difficult to treat symptoms in schizophrenia.

Our findings also justify future pioglitazone studies examining different types of memory. For example, pioglitazone may independently influence associative learning, spatial or working memory, cognitive flexibility, long-term memory, or recognition memory. Pioglitazone is also an FDA approved drug, and further human trials examining pioglitazone, antipsychotic treatment, and a larger number of specific cognitive aspects would be useful. This could be achieved through measures such as the Stroop word-color task (selective attention), n back test and backward digit span tests (working memory), Brown-Peterson test (memory capacity), or Wisconsin card sorting (planning and set-shifting)(125, 130). Other indices of cognition such as the Verbal Comprehension Index (VCI), the Perceptual Organizational Index (POI), the Working Memory Index (WMI), the Processing Speed Index (PSI), the Immediate Memory Index (IMI) and the General Memory Index are useful in human cognitive battery (125). In summary, these data suggest that pioglitazone may impact specific elements of cognition in models of schizophrenia.

### Caveats

Bioinformatics approaches have the potential to offer key insights into our understanding of how specific human diseases or healthy states manifest themselves. However, the techniques presented here have several limitations. Our pathway analyses are inherently biased as Enrichr is limited by previously published experiments, meaning some systems are more extensively studied and may skew results. Additionally, when selecting panels of clustered genes as inputs for Enrichr using dendrograms and clustering (Figure 3 and Figure 4), it is not uncommon for 5/6 of the knockdown signatures to change in the same direction for a gene, while 1/6 knockdown signatures have no change or a change in the opposite direction. However, since the goal of the study is to analyze an aggregate signature, we selected clusters of genes that largely change together in the same direction. Enrichr merges human, mouse and rat data, an approach that has advantages and disadvantages, such as comprehensive libraries but a lack of species specificity (131).

For our chemical perturbagen analyses, two seed genes (HK1 and PFKL) had the strongest discordant perturbagen signatures (Table S4), leading to the overrepresentation of these two seed genes in the overall top 20 chemical perturbagens table (Table S5). Furthermore, PFKFB2 did not have any discordant signatures to contribute. However, although not in the top 20 *overall* discordant perturbagen signatures, the top hit for PFKM was also valproic acid (albeit a weak discordance), suggesting our overall top hits may still reverse this signature.

LINCS, although a powerful resource, also has important caveats in itself. Knockdown signatures are generated in a variety of cell lines (mostly cancer) and can have differing transcriptomic changes to the same perturbagen. While the selection of a relevant cell line is important, not every target gene has been knocked down in every cell line, and comparing signatures from different cell lines can yield different results. In this study, none of our seed genes had knockdown signatures in neuronal cell lines. To minimize variability, we compared signatures for seed genes using one consistent cell line (VCAP). While these cells are not neuronal cell lines, downstream regulation of transcriptional profiles is often comparable across cell lines, and still provides a useful basis for comparison of mRNA signatures (132). This is especially true of fundamental aspects of cellular biology, including energy metabolism and related pathways considered here. Further, the LINCS database includes 978 “landmark” genes that are selected as providing representatives of classes of co-expressed genes and to enable imputing the levels of expressions for genes that are not measured directly. There may be instances where this panel does not fully capture the changes found in disease states.

Our *in silico* lookup analyses of metabolic targets in schizophrenia also present challenges. The SMRI database analysis function performs a meta-analysis across 12 independent studies, combining molecularly and functionally distinct brain regions. It is possible that for some genes, regions such as the cerebellum drive results, while more subtle differences in cortical regions are not reported. Additionally, both the SMRI and the MSSM databases were generated for whole brain regions, and are not cell-subtype specific. It is not surprising that many of our cell-subtype specific findings were not appreciated at the region-level in our lookup confirmation studies. Gene expression may be increased in certain cell types and decreased in others, with no net changes (10). The SMRI database particularly raises concern due to the number of individual studies used for comparison, many of which have varying subject demographics. This notion is supported by the observation that many of our targets are differentially regulated at the cell-level in the frontal cortical neuron database (75). Limitations aside, it will be interesting to probe these datasets (and more publicly accessible databases) for targets in pathways that were implicated in our Enrichr analyses (Table 3).

Our GluN1 knockdown work is not without limitations. While our locomotor and cognitive assay is adequately powered (n=9-12 per group), elevated plus maze, social paradigm, and PPI studies have an average of n=6 mice per group, which may not be large enough to fully elucidate subtle behavioral differences. It is also possible that pioglitazone doses used here may not fully produce therapeutic effects, and future work with varying dosing regimens is warranted. Additionally, as discussed above, cognition is multifaceted and the current study only utilizes one approach examining executive function (puzzle box assay). Several behavioral tasks such as fear conditioned learning (context versus cue), water mazes (contextual memory), radial arm maze (working memory), and novel object task would be useful in determining more nuanced effects of pioglitazone. Finally, additional biochemical experiments in GluN1 knockdown mice could help elucidate other potential metabolic pathways to target. This would provide the framework for future intervention studies using promising new pro-metabolic drugs in GluN1 knockdown mice or of pioglitazone in more models of schizophrenia (i.e. pharmacological, other mutants such as DISC1, SHANK3, NRG-1, etc.). This reverse translational approach could also generate targeted questions for future postmortem work.

In summary, we have utilized the LINCS library of genetic and chemical perturbations, coupled with a novel bioinformatics approach to gain insight into the connectivity of bioenergetic pathology in schizophrenia, generated candidate drugs with the potential to reverse the connected pathological signature as opposed to a single target, and restored long-term memory with the PPAR agonist pioglitazone in a preclinical model of schizophrenia. Our work highlights the potential for bioinformatic analyses to identify novel treatment strategies for disorders of cognition and other difficult to treat clinical domains.

## MATERIALS AND METHODS

### Human tissue acquisition and preparation

Dorsolateral prefrontal cortex (DLPFC, Brodmann area 9) postmortem brain samples originated from the Maryland Brain Collection and were distributed by both the Maryland Brain Collection and the Alabama Brain Collection. The cohort consisted of subjects with schizophrenia (n=16) and nonpsychiatrically ill comparison subjects (n=16) (Tables S1 and S2). Subjects were diagnosed with schizophrenia based on DSM-IV criteria. The medical records of the subjects were examined using a formal blinded medical chart review instrument, as well as in person interviews with the subjects and/or their caregivers, as previously described (133). Schizophrenia and comparison groups were matched for sex, age, pH, and PMI (Tables S1 and S2).

### Laser capture microdissection (LCM)

LCM was performed as previously described (11, 18, 59, 60). Briefly, 14 μm frozen tissue sections including superficial (2-3) and deep (5-6) layers of DLPFC were removed from - 80°C storage and allowed to air dry. Tissue sections were rehydrated with distilled H^2^0 and then underwent rapid Nissl staining with RNase-treated cresyl-violet (1% cresyl violet, 1% glacial acetic acid, pH 4.0). Slides were dehydrated in increasing ethanol concentrations and cleared in Histoclear II (Electron Microscopy Services, USA) for 10 minutes. After drying in a hood for 15 minutes, enriched populations of pyramidal neurons or astrocytes (1000 of each cell type per subject) were identified via morphology under the 20X objective lens and cut using the Applied Biosystems ArcturusXT™ LCM instrument and CapSure Macro LCM caps (Life Technologies, formerly Arcturus, Mountain View, CA, USA)(11, 59, 60). Laser settings were adjusted prior to each session to produce optimal cutting and capturing with laser settings ranging from 70-100 mW in power, and 2,000-3,000 μsec in duration. Separate caps were used for each subject and each cell population. Cells were captured in 4 hour sessions as we have previously demonstrated this time frame has minimal effects on messenger RNA integrity (11). Following cell capture, each cap was incubated with 50 μl of PicoPure RNA extraction buffer (Molecular Devices, Sunnyvale, CA, USA) for 30 min at 42° C. Samples were then centrifuged for 2 min at 800 × g and stored at −80°C until further processing.

### RT-qPCR

RNA was treated with RNase-free DNase (79254; Qiagen, NL) during processing. RNA was reverse-transcribed using the High-Capacity cDNA Archive Kit (Applied Biosystems, Foster City, CA, USA). Before PCR experiments, samples were preamplified as previously described (11, 59, 60).

TaqMan PCR assays for each target gene were performed in duplicate on cDNA samples in MicroAmp Fast Optical 96-well optical plates (Applied Biosystems; Foster City, CA) on an Applied Biosystems detection system (ABI SteponePlus, Applied Biosystems, Life Technologies, USA). For each reaction, 3 μl of cDNA was placed in a 20 μl reaction containing 10 μl of mastermix and 1x dilution of each primer (Applied Biosystems, Life Technologies, USA). Reactions were performed with an initial ramp time of 10 minutes at 95°C, and 40 subsequent cycles of 15 seconds at 95°C and 1 minute at 60°C. For negative controls for the qPCR reactions, cDNA was omitted (non-template control) or cDNA was generated with reverse transcriptase excluded from the reaction (no RT control). Relative concentrations of the transcripts of interest were calculated with comparison to a standard curve made with dilutions of cDNA from a pooled sampling of all the subjects. Values for the transcripts of interest were normalized to the geometric mean of B2M, β-actin (ACTB), and cyclophilin A (PPIA), housekeeping genes whose expression was unchanged in control and schizophrenia groups (Student’s t-test, p>0.05), for the same samples. All TaqMan PCR data were captured using StepOnePlus Software (StepOnePlus Real-Time PCR System; Thermoscientific, Waltham, MA)

### Primers

All primers were obtained from Thermo Fisher Scientific, Waltham, MA, USA. Human (Hs); rat (Rn). Targets are as follows: PFKL (Hs00160027_m1), GPI (Hs00976715_m1), B2M (Hs99999907_m1), ACTB (Hs99999903_m1), PPIA (Hs99999904_m1).

### Schizophrenia bioenergetic profile (seed genes)

We began by selecting genes of interest from schizophrenia pathophysiology to build a “bioenergetic schizophrenia profile.” We selected targets from the glycolytic pathway that were selectively downregulated in this study and our previous studies (GPI, HK1, PFKM, PFKL), as well as dysregulated in the literature (LDHA, PFKFB2) (Table 1)(14, 15, 18). Knockdown signatures were retrieved for each seed gene in iLINCS, and these knockdown signatures were used in subsequent lookup replication, iLINCS clustering (Enrichr), and iLINCS connectivity (drug discovery) analyses.

### *In silico* “lookup” analyses

To replicate our bioenergetic findings in schizophrenia, we probed a publicly available microarray database (Stanley Medical Research Institute Online Genomics Database), microarray data from the DLPFC of control and schizophrenia subjects (Mount Sinai School of Medicine dataset), and RNAseq data from frontal cortical neurons. The Stanley Medical Research Institute Online Genomics Database (supported by Dr. Michael Elashoff) was used to assess transcriptomic changes of metabolic targets between 50 schizophrenia and 50 control subjects in multiple brain regions (Brodmann area (BA) 6, BA8/9, BA10, BA46, and the cerebellum)(61). The Mount Sinai dataset includes microarray data from the DLPFC of schizophrenia (n=16) and control subjects (n=15). The RNAseq database contains data from frontal cortical neurons differentiated from IPSCs from a schizophrenia subject with the DISC mutation and an unaffected sibling control subject (75). We reported fold change and p-values for our metabolic targets in Table 2.

### Generation of knockdown signatures for seed genes

We used iLINCS (http://ilincs.org) to retrieve knockdown signatures for each seed gene individually. Seed genes often have multiple knockdown signatures from experiments done in different cell lines. We selected the knockdown signature from a vertebral-cancer of the prostate (VCAP) cell line for each seed gene. This was the only cell line with knockdown signatures available for all seed genes. Each seed gene knockdown signature (GPI knockdown signature, HXK1 knockdown signature, PFKM knockdown signature, PFKL knockdown signature, LDHA knockdown signature, PFKFB2 knockdown signature) is comprised of the transcriptional changes of 978 landmark genes (L1000 genes) when that seed gene is knocked down. We downloaded the knockdown signatures for each seed gene. Visual representations of the transcriptional changes in L1000 genes for each knockdown signature are presented as heatmaps generated in R Software (R version 3.4.2, 2017 The R Foundation for Statistical Computing) (Figure S3). A full list of L1000 genes is available on http://ilincs.org.

### iLINCS clustering analyses

With the goal of identifying panels of correlated transcriptomic changes in our seed gene knockdown signatures, we used iLINCS to perform unsupervised clustering of the top 50 differentially expressed L1000 genes from our 6 seed gene knockdown signatures (maximum of 300 L1000 genes) (Figure 3 and Figure 4). Due to gene overlap between top 50 differentially expressed L1000 genes in seed gene knockdown signatures, this resulted in the clustering of expression data for 260 L1000 genes. To generate this heatmap, we used the interactive “gene clusters” function in iLINCS to cluster expression data from 260 differentially expressed genes using Pearson correlation coefficients (“union of the top 50 differentially expressed genes”). Using the y-axis dendrogram function, we then selected genes that displayed similar changes in expression across the signatures. We refer to genes that are upregulated across all signatures as “panels of clustered upregulated genes” and genes that are downregulated across all signatures as “panels of clustered downregulated genes.” Panels of clustered upregulated and panels of clustered downregulated gene groups were selected separately and the data was exported to Excel (Table S3, Figure 3 and Figure 4). These clustered gene panels (Table S3) were carried forward to two separate Enrichr analyses.

### Enrichr analyses of panels of clustered upregulated and downregulated genes from iLINCS clustering analysis

To assess the biological underpinning of correlating transcriptional changes in our seed gene knockdown signatures from iLINCS, we performed traditional pathway analyses on panels of clustered upregulated and panels of clustered downregulated genes using Enrichr (http://amp.pharm.mssm.edu/Enrichr/) (Table 3, Figures S4-S17)(131, 134). Analyses for panels of clustered upregulated genes (67 genes) and panels of clustered downregulated genes (69 genes) were performed separately. We report results for KEGG cell signaling pathway, gene ontology (GO) molecular function, GO cellular components, protein-protein interacting partners, and dbGaP disease implications. Using the ChEA 2016 database in Enrichr, we also report transcription factors that have occupancy sites for a significant number of our panels of clustered genes. Finally, using LINCS L1000 database in Enrichr, we report kinases that when knocked down in specific cell lines, increase or decrease genes that are in our panels of clustered upregulated and downregulated gene data sets, respectively (Table 3, Figures S4-S17).

### iLINCS connectivity analyses (drug discovery)

With the goal of identifying small drug-like molecules with inverse signatures, we probed iLINCS for chemical perturbagens that result in L1000 transcriptomic signatures that are highly discordant (anti-correlated as denoted by negative concordance values) with each seed gene knockdown signature (potentially reversing the seed gene knockdown signature). This was done individually for each seed gene knockdown signature (unclustered) and the top 20 discordant chemical perturbagen signatures were recorded (Table S4). PFKFB2 had no discordant signatures, while GPI and PFKM had less than 20. Perturbagens appearing multiple times for the same seed gene are either in different cell lines or were administered to cells at different concentrations. We also recorded the top 20 overall discordant chemical perturbagen signatures across all seed gene knockdown signatures (Table S5). The top 20 overall discordant chemical perturbagen signatures were in the VCAP cell line. When duplicates were removed, 12 unique discordant chemical perturbagen signatures remained (Table 4). This list was use to select an FDA approved drug for preclinical trials.

### Experimental mouse lines and genotyping

Animal housing and experimentation were carried out in accordance with the Canadian Council in Animal Care (CCAC) guidelines for the care and use of animals. Mice were group housed with littermates on a 12-h light-dark cycle (0700 to 1900h) and were given access to food (2018 Teklad Global 18% Protein Rodent Diet, Envigo, Madison Wisconsin USA, www.envigo.com) ad libitum, unless otherwise specified. Mice were tail clipped and had their toes tattooed at P13 (± 3 days) for genotyping and weaned at P21. Toe tattooing was used to identify all experiment mice.

Grin1neo/neo (GluN1 knockdown) mice were generated in house, as described previously (39). The insertion mutation (neo) was identified using the following primers: wildtype forward 5’-TGA GGG GAA GCT CTT CCT GT-3’, mutant forward 5’-GCT TCC TCG TGC TTT ACG GTA T-3’, common reverse 5’-AAG CGA TTA GAC AAC TAA GGG T-3’.

For our LCMS/IPA and qPCR studies, we had 5 GluN1 knockdown and 5 WT mice. For our open field test and puzzle box assay pioglitazone studies, we had a total of 43 animals, 22 WT and 21 GluN1 knockdown mice (Table S6). For our elevated plus maze, social paradigm, and PPI pioglitazone studies, we had a total of 27 animals, 14 WT and 13 GluN1 knockdown mice (Table S6).

### GluN1 knockdown LCMS and RT-qPCR studies

We used a mouse anti-PSD-95 antibody (Millipore, catalogue # MAB1596) to capture PSD-95 protein complexes from samples (3 male WT, 3 male GluN1 knockdown). We verified the specificity of this antibody using multiple reaction monitoring mass spectrometry analysis of PSD-95 peptides captured by affinity purification. 5ug of PSD-95 antibody was coupled per 1mg of Dynabeads (Life Technologies) according to the antibody coupling kit protocol (#14311D). For each sample, 1000ul of 10mg/ml antibody coupled beads were washed 2x 1ml with ice cold 1x PBST (#9809S, Cell Signaling) then incubated with mouse brain lysate brought to a final volume of 1000ul with ice cold 1x PBST for 1 hour at room temperature. The supernatant was removed and the beads were washed 4 × 10 minutes at room temperature in 1ml ice cold 1x PBST. Captured protein complexes were eluted with 30ul of 1N Ammonium Hydroxide (#320145, Sigma), 5mM EDTA, pH 12 for 10 minutes at room temperature. 6ul of 6x protein denaturing buffer (4.5% SDS, 15% β-mercaptoethanol, 0.018% bromophenol blue, and 36% glycerol in 170 mM Tris-HCl, pH 6.8) was added to each sample elution. The eluted samples were heated at 70°C for 10 minutes then processed for mass spectrometry.

All samples (3 male WT, 3 male GluN1 knockdown) were loaded on a 1.5 mm, 4-12% Bis-Tris Invitrogen NuPage gel (NP0335BOX) and electrophoresed in 1x MES buffer (NP0002) for 10 minutes at 180v. The gel was fixed in 50% ethanol/10% acetic acid overnight at RT, then washed in 30% ethanol for 10 min followed by two 10 min washes in MilliQ water (MilliQ Gradient system). The lanes were harvested, cut into small (∼2mm) squares, and subjected to in-gel tryptic digestion and peptide recovery. Representative western blots are seen in Figure 5A. Samples were resuspended in 0.1% formic acid.

Nano liquid chromatography coupled electrospray tandem mass spectrometry (nLC-ESI-MS/MS) analyses were performed on a 5600+ QTOF mass spectrometer (Sciex, Toronto, On, Canada) interfaced to an Eksigent (Dublin, CA) nanoLC.ultra nanoflow system. Peptides were loaded (via an Eksigent nanoLC.as-2 autosampler) onto an IntegraFrit Trap Column (outer diameter of 360 μm, inner diameter of 100, and 25 μm packed bed) from New Objective, Inc. (Woburn, MA) at 2 μl/min in formic acid/H_2_O 0.1/99.9 (v/v) for 15 min to desalt and concentrate the samples. For the chromatographic separation of peptides, the trap-column was switched to align with the analytical column, Acclaim PepMap100 (inner diameter of 75 μm, length of 15 cm, C18 particle sizes of 3 μm and pore sizes of 100 Å) from Dionex-Thermo Fisher Scientific (Sunnyvale, CA). The peptides were eluted using a variable mobile phase (MP) gradient from 95% phase A (Formic acid/H_2_O 0.1/99.9, v/v) to 40% phase B (Formic Acid/Acetonitrile 0.1/99.9, v/v) for 70 min, from 40% phase B to 85% phase B for 5 min and then keeping the same mobile phase composition for 5 additional min at 300 nL/min. The nLC effluent was ionized and sprayed into the mass spectrometer using NANOSpray^®^ III Source (Sciex). Ion source gas 1 (GS1), ion source gas 2 (GS2) and curtain gas (CUR) were respectively kept at 8, 0 and 35 vendor specified arbitrary units. The mass spectrometer method was operated in positive ion mode and the interface heater temperature and ion spray voltage were kept at 150°C, and at 2.6 kV, respectively. The data was recorded using Analyst-TF (version 1.7) software.

The data independent acquisition (DIA) method was set to go through 1757 cycles for 99 minutes, where each cycle performed one TOF-MS scan type (0.25 sec accumulation time, in a 550.0 to 830.0 m/z window) followed by 56 sequential overlapping windows of 6 Daltons each. Note that the Analyst software automatically adds 1 Dalton to each window to provide overlap, thus an input of 5 Da in the method set up window results in an overlapping 6 Da collection window width (e.g. 550-556, then 555-561, 560-566, etc). Within each window, a charge state of +2, high sensitivity mode, and rolling collision energy with a collision energy spread (CES) of 15 V was selected.

Protalizer DIA software (Vulcan Analytical, Birmingham, AL) was used to analyze every DIA file (Sciex 5600 QTOFs). The Swiss-Prot *Homo Sapien* database, downloaded March 17^th^ 2015, was used as the reference database for all MS/MS searches. A precursor and fragment-ion tolerance for QTOF instrumentation was used for the Protalizer Caterpillar spectral-library free identification algorithm. Potential modifications included in the searches were phosphorylation at S, T, and Y residues, N-terminal acetylation, N-terminal loss of ammonia at C residues, and pyroglutamic acid at N-terminal E and Q residues. Carbamidomethylation of C residues was searched as a fixed modification. The maximum valid protein and peptide expectation score from the X! Tandem search engine used for peptide and protein identification on reconstructed spectra was set to 0.005.

For DIA quantification by Protalizer the maximum number of b and y series fragment-ion transitions were set to nine excluding those with *m/z* values below 300 and not containing at least 10% of the relative intensity of the strongest fragment-ion assigned to a peptide. A minimum of five fragment-ions were required for a peptide to be quantified (except where indicated otherwise). In datasets where a minimum of seven consistent fragment-ions were not detected for the same peptide ion in each of the three files compared in a triplicate analysis, the algorithm identified the file with the largest sum fragment-ion AUC and extracted up to seven of these in the other files using normalized retention time coordinates based on peptides detected by the Caterpillar algorithm in all the files in a dataset.

To generate the top pathways for GluN1 knockdown mice we input the top 20 increased proteins in GluN1 knockdown mice relative to WT mice from LCMS experiments into Ingenuity Pathway Analysis (IPA). The top 5 implicated pathways are reported in Figure 5B.

Finally, we examined the mRNA expression of metabolic transporters using real time quantitative polymerase chain reaction (RT-qPCR) in a cohort of GluN1 knockdown and wildtype (WT) mice (n=5 per group) (Figure 5C). TaqMan PCR assays for each target gene were performed in duplicate on cDNA samples in 96-well optical plates using StepOnePlus machine (StepOnePlus Real-Time PCR System; Thermoscientific, Waltham, MA). All TaqMan PCR data were captured using Sequence Detector Software (SDS version 1.6; PE Applied Biosystems). All primers were obtained from Thermo Fisher Scientific, Waltham, MA, USA. Targets are as follows: MCT1 (Mm01306379_m1), MCT2 (Mm00441442_m1), MCT4 (Mm00446102_m1), GLUT1 (Mm00441480_m1), and GLUT3 (Mm00441483_m1).

### Drug administration for GluN1 knockdown pioglitazone studies

Pioglitazone is an FDA approved drug for the treatment of type II diabetes. It increases the transport of glucose from the bloodstream into tissues by increasing the expression of the glucose transporter GLUT1. We divided mice into 4 groups: for the open field test and puzzle box assay, WTveh (n=12), WTpio (n=10), GluN1veh (n=9) and GluN1pio (n=12) and for elevated plus maze, social paradigm, and pre-pulse inhibition tasks, WTveh (n=6), WTpio (n=8), GluN1veh (n=4) and GluN1pio (n=9). On day 1 of the experimental paradigm, mice were given free access to either a 2018 Teklad Global 18% Protein Rodent Diet (Envigo, Madison Wisconsin USA, www.envigo.com) as control chow or chow infused with pioglitazone at 100 ppm (2018 Teklad with 0.01% pio, BOCSCI, Shirley, NY, USA) for 7 days. Dose was selected based on previous studies (54). On day 8, behavioral tests began (animals remained on respective diets until sacrificed on day 17). Throughout the experiments, we assessed caloric intake and body weight to ensure pioglitazone treatment did not have adverse effects (Figure S18).

### Behavioral testing

F1 male and female Grin1fl^neo/Cre^ mice were used for all behavioral testing, along with WT littermates as controls. Animals were aged 10-12 weeks. All behavioral tests were completed between 09:00 and 15:00h. Animals were weighed at the beginning of the experiment and on Days 7 and 14. Food was also weighed throughout experiments (Figure S18). On Day 7, all mice had their locomotor activity and stereotypy behavior measured in the Open Field Test. On day 9, mice were tested on the Elevated Plus Maze (EPM). Days 10 through 12, mice were tested in the puzzle box assay. Day 15 mice were subjected to the social affiliative paradigm, and finally on day 16, pre-pulse inhibition was tested. A timetable is presented in Table S7.

### Open Field Test

Locomotor activity and stereotypy were measured using digital activity monitors (Omnitech Electronics, Columbus, OH, USA) on the first day of behavioral testing (Day 8 of overall experimental paradigm) as previously described (135). Naïve mice were placed in novel Plexiglas arenas (20 × 20 × 45 cm) and their locomotor and stereotypic activity were recorded over a 120-min period in dim light (15-16lux). Activity was tracked via infrared light beam sensors; total distance traveled and stereotypic movements were collected in 5-min bins.

### Elevated Plus Maze

Anxiety behavior was assessed on day 9 of testing via the elevated plus maze (136). The elevated plus maze was composed of 4 opaque-white arms (2 opposite arms closed, 2 opposite arms open), arranged in a plus shape, with an open center. The dimensions were as follows; maze elevation (38.7cm), open arm (L:30.5cm, W:5cm, H:0cm), closed arm (L:30.5cm, W:5cm, H:15.2cm) and center (5cm x 5cm). The experiment mouse was placed in the center of the maze, and allowed to freely explore the maze for 8 min. in dim light (15-16lux), while being tracked with an overhead camera. Open and closed arm times were recorded and collected by Biobserve Viewer3 software. The percentage of time spent in the open arms, as compared to closed arms and center time, was calculated and expressed as a percent of time spent in the open arms of the maze.

### Puzzle Box Assay

Mice were run on the Puzzle Box Assay on days 10-12 of testing to assess cognition and executive function as previously described (135). Consisting of two compartments, the puzzle box contains a start area (58 × 28 × 27.5 cm) in bright light (250lux), and a goal zone (14 × 28 × 27.5 cm) in dim light (5lux). The two areas are separated by a black Plexiglas divider, but connected via an underpass large enough for mice to pass through easily. Mice were placed in the start box facing away from the divider, and the time to move to the goal zone (through the underpass, with both hind legs in the goal zone) was manually scored and recorded. Mice were tested over three days, 3 trials/day, with each day consisting of increasingly difficult obstacles present in the underpass connecting the start area and the goal zone. 2 min were given between each trial on a given day, with a max 300 sec allowed for the completion of each trial. The trials (and obstacles) for the puzzle box were as follows:

*Day 10*: T1 (training) open door and unblocked underpass, T2 and T3 (challenge, then learning) doorway closed and underpass open
*Day 11*: T4 (long-term memory) identical to T3, T5 and T6 (challenge, then learning) underpass filled with bedding (similar to that found in home cage)
*Day 12*: T7 (long-term memory) identical to T6, T8 and T9 (challenge, then learning) underpass blocked by a removable cardboard plug

### Social Affiliative Paradigm

Social affiliative behavior was assessed on day 15 of testing, as previously described (135, 137). Sociability was measured via video recording the motion and exploration of the experimental mouse, tracked via Biobserve Viewer (version 2) software (center body – reference point). Experimental mice were allowed to explore the open area (opaque white walls, 62 × 42 × 22 cm) for 10-min. in dim lighting (15-16lux). The area contained two inverted wire cups, one containing a stimulus mouse (‘social’) and the other empty (‘non-social’). Time spent in each zone (3 cm zone around the cup) was recorded via the Biobserve software. Mice used as a social stimulus were novel, wildtype, inbred C57Bl/6 mice that were age-and sex-matched to the test mouse.

### Pre-Pulse Inhibition/Acoustic Startle Reflex

Pre-pulse inhibition of the acoustic startle response was measured on day 16, via SR-LAB equipment and software from San Diego Instruments (San Diego Instruments, San Diego, CA, USA). Accelerometers were calibrated to 700±5 mV and output voltages were amplified and analyzed for voltage changes using SR Analysis, and exported as an excel file. Background white noise was maintained at 65dB. PPI was measured in a 30-min test with 80 randomized trials of: (1) 10 trials pulse alone (2) 10 trials pre-pulse alone (for each pre-pulse), (3) 10 trials pre-pulse plus pulse (for each pre-pulse), and (4) 10 trials no pulse. 5 pulse alone trials were performed before and after the 80 trials, totaling 90 trials per run. The pre-pulse (4dB, 8dB, or 16dB) was presented 100ms prior to the startle pulse (165dB). The inter-stimulus interval (ISI) was randomized between 5 and 20s. Experimental mice were placed in a cylindrical tube on a platform in a soundproof chamber. Mice were allowed to acclimatize in the chamber and to the background noise for 300s, followed by 5 consecutive pulse alone trials, then by 80 randomized trials (as described above) and then 5 consecutive pulse alone trials. Pre-pulse inhibition was measured as a decrease in the amplitude of startle response to a 100dB acoustic startle pulse, following each pre-pulse (4dB, 8dB and 16dB).

### Statistical analysis

Statistical analyses for human LCM-qPCR were as follows: data were analyzed using Statistica 13.0 (Statsoft, Tulsa, Oklahoma, USA) and GraphPad Prism (La Jolla, CA, USA). Outliers were removed from data sets using the ROUT method (Q = 5%). All dependent measures were tested for normalcy and homogeneity of variance using D'Agostino & Pearson omnibus normality test followed by Student’s t-tests. We probed for associations between our dependent measures and pH, PMI, age, and RIN using correlation analyses. We did not detect any significant correlations between age, pH, and PMI with any of our dependent measures.

In addition to postmortem interval, pH, and age, we performed secondary analyses (using t-tests) to examine ethanol use, smoking, and laterality as grouping variables in our schizophrenia subjects and found no significant effects. We also performed t-tests after removing antipsychotic naïve subjects and all targets remained significant. No subjects had any dependence for cocaine, sedatives, cannabis, amphetamines, hallucinogens, inhalants, or opioid dependence (only subject 828 took morphine for pain). Subjects with prolonged agonal states were not included in this cohort.

Statistical analyses for animal behavioral assays were as follows: All dependent measures were tested for normalcy and homogeneity of variance using D'Agostino & Pearson omnibus normality test. For locomotor activity, stereotypy, vertical activity, EPM, social and social novelty, and PPI/acoustic startle response, we performed 2-way ANOVA with Bonferonni multiple comparison corrections. Each dB in PPI was treated as an individual ANOVA (4 dB, 8 dB, 16 dB).

## ACKNOWLEDGEMENTS

This work was funded by the following grants: MH107487, MH107916, Local Initiative for Excellence (L.I.F.E.) Foundation, MH107487, ES006096, HL127624, UL1TR001425, and CIHR Operating Grant (119298).

## Data and availability

The Mount Sinai School of Medicine microarray data from this publication can be accessed at https://harouv01.u.hpc.mssm.edu/ (PMID 22868662). The Stanley Medical Research Institute genomics database is publically available at https://www.stanleygenomics.org/stanley/stat.jsp. The IPSC RNAseq data were deposited at GEO (accession number: GSE57821).

## AUTHOR CONTRIBUTIONS

CS, JM, and RM designed the study. JM, ED, EB, SO, AF, CS, and RM developed the bioinformatic workflow. CS performed human, *in silico*, and iLINCS analyses. AR provided the GluN1 knockdown mouse model. AF performed GluN1 knockdown immunoprecipitation and LCMS studies. CM performed all GluN1 knockdown behavior studies. PK, MP, ZW, and VH provided lookup study databases. All authors read and approved the final manuscript.

## CONFLICT OF INTEREST

The authors have no conflict of interest to declare.

## REFERENCES

1. Association AP. Diagnostic and Statistical Manual of Mental Disorders. Fourth, Text Revision ed. Washington, D.C.: American Psychiatric Association; 2000.

2. Buchanan RW, Carpenter WT. Schizophrenia: Introduction and overview. In: Sadock BJ, Sadock VA, editors. Comprehensive Textbook of Psychiatry. 1. Philadelphia: Lippincott, Williams, and Wilkins; 2000. p. 1096–110.

3. Fleischhacker W. Negative symptoms in patients with schizophrenia with special reference to the primary versus secondary distinction. L’Encephale. 2000;26 Spec No 1:12–4.

4. Zanello A, Curtis L, Badan Ba M, Merlo MC. Working memory impairments in first-episode psychosis and chronic schizophrenia. Psychiatry research. 2009;165(1-2):10–8.

5. Potkin SG, Turner JA, Brown GG, McCarthy G, Greve DN, Glover GH, et al. Working memory and DLPFC inefficiency in schizophrenia: the FBIRN study. Schizophrenia bulletin. 2009;35(1):19–31.

6. Wobrock T, Schneider M, Kadovic D, Schneider-Axmann T, Ecker UK, Retz W, et al. Reduced cortical inhibition in first-episode schizophrenia. Schizophrenia research. 2008;105(1-3):252–61.

7. Meltzer HY, McGurk SR. The effects of clozapine, risperidone, and olanzapine on cognitive function in schizophrenia. Schizophrenia bulletin. 1999;25(2):233–55.

8. Meltzer HY, Park S, Kessler R. Cognition, schizophrenia, and the atypical antipsychotic drugs. Proceedings of the National Academy of Sciences. 1999;96(24):13591.

9. Goldman-Rakic PS. Cellular basis of working memory. Neuron. 1995;14(3):477–85.

10. McCullumsmith RE, Hammond JH, Shan D, Meador-Woodruff JH. Postmortem brain: an underutilized substrate for studying severe mental illness. Neuropsychopharmacology: official publication of the American College of Neuropsychopharmacology. 2014;39(1):65–87.

11. McCullumsmith RE, O'Donovan SM, Drummond JB, Benesh FS, Simmons M, Roberts R, et al. Cell-specific abnormalities of glutamate transporters in schizophrenia: sick astrocytes and compensating relay neurons? Mol Psychiatry. 2016;6:823–30.

12. McCullumsmith R, Clinton S, Meador-Woodruff J. Schizophrenia as a disorder of neuroplasticity. International review of neurobiology. 2004;59:19–45.

13. Sullivan CR, O’Donovan S, McCullumsmith RE, Ramsey A. Defects in bioenergetic coupling in schizophrenia. Biological psychiatry. 2017.

14. Stone WS, Faraone SV, Su J, Tarbox SI, Van Eerdewegh P, Tsuang MT. Evidence for linkage between regulatory enzymes in glycolysis and schizophrenia in a multiplex sample. American journal of medical genetics Part B, Neuropsychiatric genetics: the official publication of the International Society of Psychiatric Genetics. 2004;127B(1):5–10.

15. Altar CA, Jurata LW, Charles V, Lemire A, Liu P, Bukhman Y, et al. Deficient hippocampal neuron expression of proteasome, ubiquitin, and mitochondrial genes in multiple schizophrenia cohorts. Biological psychiatry. 2005;58(2):85–96.

16. Arion D, Corradi JP, Tang S, Datta D, Boothe F, He A, et al. Distinctive transcriptome alterations of prefrontal pyramidal neurons in schizophrenia and schizoaffective disorder. Mol Psychiatry. 2015;20(11):1397–405.

17. Arion D, Huo Z, Enwright JF, Corradi JP, Tseng G, Lewis DA. Transcriptome alterations in prefrontal pyramidal cells distinguish schizophrenia from bipolar and major depressive disorders. Biological psychiatry.

18. Courtney Sullivan RK, Kathryn Hasselfeld, Sinead O'Donovan, Amy Ramsey, Robert McCullumsmith. Neuron-specific deficits of neuroenergetic processes in the dorsolateral prefrontal cortex in schizophrenia. Molecular Psychiatry. 2018;in press.

19. Koleti A, Terryn R, Stathias V, Chung C, Cooper DJ, Turner JP, et al. Data Portal for the Library of Integrated Network-based Cellular Signatures (LINCS) program: integrated access to diverse large-scale cellular perturbation response data. Nucleic acids research. 2018;46(D1):D558–d66.

20. Duan Q, Flynn C, Niepel M, Hafner M, Muhlich JL, Fernandez NF, et al. LINCS Canvas Browser: interactive web app to query, browse and interrogate LINCS L1000 gene expression signatures. Nucleic acids research. 2014;42(Web Server issue):W449–W60.

21. Actos (Pioglitazone Hydrochloride). Takeda Armerica Research & Dev. Ctr. Inc.. Deerfield, IL1999.

22. Jentsch JD, Roth RH. The neuropsychopharmacology of phencyclidine: from NMDA receptor hypofunction to the dopamine hypothesis of schizophrenia. Neuropsychopharmacology: official publication of the American College of Neuropsychopharmacology. 1999;20(3):201–25.

23. Allen R, Young S. Phencyclidine-induced psychosis. The American journal of psychiatry. 1978;135(9):1081–4.

24. Javitt DC, Zukin SR. Recent advances in the phencyclidine model of schizophrenia. The American journal of psychiatry. 1991;148(10):1301–8.

25. Coyle JT, Tsai G, Goff DC. Ionotropic glutamate receptors as therapeutic targets in schizophrenia. Current drug targets CNS and neurological disorders. 2002;1(2):183–9.

26. Luby ED, Cohen BD, Rosenbaum G, Gottlieb JS, Kelley R. Study of a new schizophrenomimetic drug; sernyl. AMA archives of neurology and psychiatry. 1959;81(3):363–9.

27. Krystal JH, Karper LP, Seibyl JP, Freeman GK, Delaney R, Bremner JD, et al. Subanesthetic effects of the noncompetitive NMDA antagonist, ketamine, in humans. Psychotomimetic, perceptual, cognitive, and neuroendocrine responses. Archives of general psychiatry. 1994;51(3):199–214.

28. Lahti AC, Weiler MA, Tamara Michaelidis BA, Parwani A, Tamminga CA. Effects of ketamine in normal and schizophrenic volunteers. Neuropsychopharmacology: official publication of the American College of Neuropsychopharmacology. 2001;25(4):455–67.

29. Kantrowitz JT, Javitt DC. N-methyl-d-aspartate (NMDA) receptor dysfunction or dysregulation: the final common pathway on the road to schizophrenia? Brain research bulletin. 2010;83(3-4):108–21.

30. Begni S, Moraschi S, Bignotti S, Fumagalli F, Rillosi L, Perez J, et al. Association between the G1001C polymorphism in the GRIN1 gene promoter region and schizophrenia. Biological psychiatry. 2003;53(7):617–9.

31. Georgi A, Jamra RA, Klein K, Villela AW, Schumacher J, Becker T, et al. Possible association between genetic variants at the GRIN1 gene and schizophrenia with lifetime history of depressive symptoms in a German sample. Psychiatric genetics. 2007;17(5):308–10.

32. Galehdari H, Pooryasin A, Foroughmand A, Daneshmand S, Saadat M. Association between the G1001C polymorphism in the GRIN1 gene promoter and schizophrenia in the Iranian population. Journal of molecular neuroscience: MN. 2009;38(2):178–81.

33. Itokawa M, Yamada K, Yoshitsugu K, Toyota T, Suga T, Ohba H, et al. A microsatellite repeat in the promoter of the N-methyl-D-aspartate receptor 2A subunit (GRIN2A) gene suppresses transcriptional activity and correlates with chronic outcome in schizophrenia. Pharmacogenetics. 2003;13(5):271–8.

34. Iwayama-Shigeno Y, Yamada K, Itokawa M, Toyota T, Meerabux JM, Minabe Y, et al. Extended analyses support the association of a functional (GT)n polymorphism in the GRIN2A promoter with Japanese schizophrenia. Neuroscience letters. 2005;378(2):102–5.

35. Tang J, Chen X, Xu X, Wu R, Zhao J, Hu Z, et al. Significant linkage and association between a functional (GT)n polymorphism in promoter of the N-methyl-D-aspartate receptor subunit gene (GRIN2A) and schizophrenia. Neuroscience letters. 2006;409(1):80–2.

36. Qin S, Zhao X, Pan Y, Liu J, Feng G, Fu J, et al. An association study of the N-methyl-D-aspartate receptor NR1 subunit gene (GRIN1) and NR2B subunit gene (GRIN2B) in schizophrenia with universal DNA microarray. European journal of human genetics: EJHG. 2005;13(7):807–14.

37. Martucci L, Wong AH, De Luca V, Likhodi O, Wong GW, King N, et al. N-methyl-D-aspartate receptor NR2B subunit gene GRIN2B in schizophrenia and bipolar disorder: Polymorphisms and mRNA levels. Schizophrenia research. 2006;84(2-3):214–21.

38. Demontis D, Nyegaard M, Buttenschon HN, Hedemand A, Pedersen CB, Grove J, et al. Association of GRIN1 and GRIN2A-D with schizophrenia and genetic interaction with maternal herpes simplex virus-2 infection affecting disease risk. American journal of medical genetics Part B, Neuropsychiatric genetics: the official publication of the International Society of Psychiatric Genetics. 2011;156b(8):913–22.

39. Mohn AR, Gainetdinov RR, Caron MG, Koller BH. Mice with reduced NMDA receptor expression display behaviors related to schizophrenia. Cell. 1999;98(4):427–36.

40. Ramsey AJ. NR1 knockdown mice as a representative model of the glutamate hypothesis of schizophrenia. Progress in brain research. 2009;179:51–8.

41. Gandal MJ, Sisti J, Klook K, Ortinski PI, Leitman V, Liang Y, et al. GABAB-mediated rescue of altered excitatory-inhibitory balance, gamma synchrony and behavioral deficits following constitutive NMDAR-hypofunction. Translational psychiatry. 2012;2:e142.

42. Wesseling H, Guest PC, Lee CM, Wong EH, Rahmoune H, Bahn S. Integrative proteomic analysis of the NMDA NR1 knockdown mouse model reveals effects on central and peripheral pathways associated with schizophrenia and autism spectrum disorders. Molecular autism. 2014;5:38.

43. Zhou K, Yang Y, Gao L, He G, Li W, Tang K, et al. NMDA receptor hypofunction induces dysfunctions of energy metabolism and semaphorin signaling in rats: a synaptic proteome study. Schizophrenia bulletin. 2012;38(3):579–91.

44. Sun L, Li J, Zhou K, Zhang M, Yang J, Li Y, et al. Metabolomic analysis reveals metabolic disturbance in the cortex and hippocampus of subchronic MK-801 treated rats. PloS one. 2013;8(4):e60598.

45. Steinman MQ, Gao V, Alberini CM. The Role of Lactate-Mediated Metabolic Coupling between Astrocytes and Neurons in Long-Term Memory Formation. Frontiers in Integrative Neuroscience. 2016;10:10.

46. Suzuki A, Stern SA, Bozdagi O, Huntley GW, Walker RH, Magistretti PJ, et al. Astrocyte-neuron lactate transport is required for long-term memory formation. Cell. 2011;144(5):810–23.

47. Newman LA, Korol DL, Gold PE. Lactate produced by glycogenolysis in astrocytes regulates memory processing. PloS one. 2011;6(12):e28427.

48. Smith U. Pioglitazone: mechanism of action. International journal of clinical practice Supplement. 2001(121):13–8.

49. Seto SW, Yan Lam T, Leung G, L.S. Au A, Ngai SM, Wan Chan S, et al. Comparison of vascular relaxation, lipolysis and glucose uptake by peroxisome proliferator-activated receptor-γ activation in + db/+ m and + db/+ db mice 2007. 40–8 p.

50. Colca JR, McDonald WG, Waldon DJ, Leone JW, Lull JM, Bannow CA, et al. Identification of a novel mitochondrial protein ("mitoNEET") cross-linked specifically by a thiazolidinedione photoprobe. Am J Physiol Endocrinol Metab. 2004;286(2):E252–60.

51. Paddock ML, Wiley SE, Axelrod HL, Cohen AE, Roy M, Abresch EC, et al. MitoNEET is a uniquely folded 2Fe 2S outer mitochondrial membrane protein stabilized by pioglitazone. Proceedings of the National Academy of Sciences of the United States of America. 2007;104(36):14342–7.

52. Hwang J, Kleinhenz DJ, Rupnow HL, Campbell AG, Thule PM, Sutliff RL, et al. The PPARgamma ligand, rosiglitazone, reduces vascular oxidative stress and NADPH oxidase expression in diabetic mice. Vascular pharmacology. 2007;46(6):456–62.

53. Wright MB, Bortolini M, Tadayyon M, Bopst M. Minireview: Challenges and Opportunities in Development of PPAR Agonists. Molecular Endocrinology. 2014;28(11):1756–68.

54. Dello Russo C, Gavrilyuk V, Weinberg G, Almeida A, Bolanos JP, Palmer J, et al. Peroxisome proliferator-activated receptor gamma thiazolidinedione agonists increase glucose metabolism in astrocytes. The Journal of biological chemistry. 2003;278(8):5828–36.

55. Amann LC, Gandal MJ, Halene TB, Ehrlichman RS, White SL, McCarren HS, et al. Mouse behavioral endophenotypes for schizophrenia. Brain research bulletin. 2010;83(3):147–61.

56. Ben Abdallah NM, Fuss J, Trusel M, Galsworthy MJ, Bobsin K, Colacicco G, et al. The puzzle box as a simple and efficient behavioral test for exploring impairments of general cognition and executive functions in mouse models of schizophrenia. Exp Neurol. 2011;227(1):42–52.

57. Ramsey AJ, Milenkovic M, Oliveira AF, Escobedo-Lozoya Y, Seshadri S, Salahpour A, et al. Impaired NMDA receptor transmission alters striatal synapses and DISC1 protein in an age-dependent manner. Proc Natl Acad Sci U S A. 2011;108(14):5795–800.

58. Geyer MA, McIlwain KL, Paylor R. Mouse genetic models for prepulse inhibition: an early review. Mol Psychiatry. 2002;7(10):1039–53.

59. O'Donovan SM, Hasselfeld K, Bauer D, Simmons M, Roussos P, Haroutunian V, et al. Glutamate transporter splice variant expression in an enriched pyramidal cell population in schizophrenia. Translational psychiatry. 2015;5:e579.

60. Sodhi MS, Simmons M, McCullumsmith R, Haroutunian V, Meador-Woodruff JH. Glutamatergic gene expression is specifically reduced in thalamocortical projecting relay neurons in schizophrenia. Biological psychiatry. 2011;70(7):646–54.

61. Higgs BW, Elashoff M, Richman S, Barci B. An online database for brain disease research. BMC Genomics. 2006;7:70-.

62. Gross DN, Wan M, Birnbaum MJ. The role of FOXO in the regulation of metabolism. Current diabetes reports. 2009;9(3):208–14.

63. Wang Y, Zhou Y, Graves DT. FOXO Transcription Factors: Their Clinical Significance and Regulation. BioMed research international. 2014;2014:925350.

64. Gehart H, Kumpf S, Ittner A, Ricci R. MAPK signalling in cellular metabolism: stress or wellness? EMBO Reports. 2010;11(11):834–40.

65. Sethi JK, Vidal-Puig A. Wnt signalling and the control of cellular metabolism. The Biochemical journal. 2010;427(1):1–17.

66. Iranpour N, Zandifar A, Farokhnia M, Goguol A, Yekehtaz H, Khodaie-Ardakani MR, et al. The effects of pioglitazone adjuvant therapy on negative symptoms of patients with chronic schizophrenia: a double-blind and placebo-controlled trial. Human psychopharmacology. 2016;31(2):103–12.

67. Koster JF, Slee RG, Van Berkel TJ. Isoenzymes of human phosphofructokinase. Clinica chimica acta; international journal of clinical chemistry. 1980;103(2):169–73.

68. Van Schaftingen E, Hue L, Hers HG. Fructose 2,6-bisphosphate, the probably structure of the glucose- and glucagon-sensitive stimulator of phosphofructokinase. Biochem J. 1980;192(3):897–901.

69. Graham DB, Becker CE, Doan A, Goel G, Villablanca EJ, Knights D, et al. Functional genomics identifies negative regulatory nodes controlling phagocyte oxidative burst. Nature Communications. 2015;6:7838.

70. Annerén KG, Korenberg JR, Epstein CJ. Phosphofructokinase activity in fibroblasts aneuploid for chromosome 21. Human Genetics. 1987;76(1):63–5.

71. Chaput M, Claes V, Portetelle D, Cludts I, Cravador A, Burny A, et al. The neurotrophic factor neuroleukin is 90% homologous with phosphohexose isomerase. Nature. 1988;332(6163):454–5.

72. Liu M-L, Zhang X-T, Du X-Y, Fang Z, Liu Z, Xu Y, et al. Severe disturbance of glucose metabolism in peripheral blood mononuclear cells of schizophrenia patients: a targeted metabolomic study. Journal of Translational Medicine. 2015;13:226.

73. Haller JF, Krawczyk SA, Gostilovitch L, Corkey BE, Zoeller RA. Glucose-6-phosphate isomerase deficiency results in mTOR activation, failed translocation of lipin 1α to the nucleus and hypersensitivity to glucose: Implications for the inherited glycolytic disease. Biochimica et biophysica acta. 2011;1812(11):1393–402.

74. Gururajan A, van den Buuse M. Is the mTOR-signalling cascade disrupted in Schizophrenia? Journal of neurochemistry. 2014;129(3):377–87.

75. Wen Z, Nguyen HN, Guo Z, Lalli MA, Wang X, Su Y, et al. Synaptic dysregulation in a human iPS cell model of mental disorders. Nature. 2014;515(7527):414–8.

76. Nicholson KM, Anderson NG. The protein kinase B/Akt signalling pathway in human malignancy. Cellular signalling. 2002;14(5):381–95.

77. Guan X-M, Yu H, Jiang Q, Van der Ploeg LHT, Liu Q. Distribution of neuromedin U receptor subtype 2 mRNA in the rat brain☆☆Published on the World Wide Web on 27 November 2000. Gene Expression Patterns. 2001;1(1):1–4.

78. Nakazato M, Hanada R, Murakami N, Date Y, Mondal MS, Kojima M, et al. Central Effects of Neuromedin U in the Regulation of Energy Homeostasis. Biochemical and Biophysical Research Communications. 2000;277(1):191–4.

79. Pal R, Janz M, Galson DL, Gries M, Li S, Johrens K, et al. C/EBPbeta regulates transcription factors critical for proliferation and survival of multiple myeloma cells. Blood. 2009;114(18):3890–8.

80. Ruffell D, Mourkioti F, Gambardella A, Kirstetter P, Lopez RG, Rosenthal N, et al. A CREB-C/EBPβ cascade induces M2 macrophage-specific gene expression and promotes muscle injury repair. Proceedings of the National Academy of Sciences of the United States of America. 2009;106(41):17475–80.

81. Kelly RD, Cowley SM. The physiological roles of histone deacetylase (HDAC) 1 and 2: complex costars with multiple leading parts. Biochemical Society transactions. 2013;41(3):741–9.

82. Moldovan G-L, Pfander B, Jentsch S. PCNA, the Maestro of the Replication Fork. Cell. 129(4):665–79.

83. Jakovcevski M, Bharadwaj R, Straubhaar J, Gao G, Gavin DP, Jakovcevski I, et al. Prefrontal cortical dysfunction after overexpression of histone deacetylase 1. Biological psychiatry. 2013;74(9):696–705.

84. Narayan S, Tang B, Head SR, Gilmartin TJ, Sutcliffe JG, Dean B, et al. Molecular Profiles of Schizophrenia in the CNS at Different Stages of Illness. Brain research. 2008;1239:235–48.

85. Sharma RP, Grayson DR, Gavin DP. Histone deactylase 1 expression is increased in the prefrontal cortex of schizophrenia subjects: analysis of the National Brain Databank microarray collection. Schizophrenia research. 2008;98(1-3):111–7.

86. Akbarian S. Epigenetic mechanisms in schizophrenia. Dialogues in Clinical Neuroscience. 2014;16(3):405–17.

87. Chandra V, Huang P, Hamuro Y, Raghuram S, Wang Y, Burris TP, et al. Structure of the intact PPAR-γ–RXR-α nuclear receptor complex on DNA. Nature. 2008;456(7220):350–6.

88. Lehmann JM, Moore LB, Smith-Oliver TA, Wilkison WO, Willson TM, Kliewer SA. An Antidiabetic Thiazolidinedione Is a High Affinity Ligand for Peroxisome Proliferator-activated Receptor γ (PPARγ). Journal of Biological Chemistry. 1995;270(22):12953–6.

89. Muller N. Inflammation and the glutamate system in schizophrenia: implications for therapeutic targets and drug development. Expert Opin Ther Targets. 2008;12(12):1497–507.

90. Muller N, Myint AM, Schwarz MJ. Inflammation in schizophrenia. Adv Protein Chem Struct Biol. 2012;88:49–68.

91. Müller N, Schwarz MJ. Immune System and Schizophrenia. Current immunology reviews. 2010;6(3):213–20.

92. Gandal MJ, Haney JR, Parikshak NN, Leppa V, Ramaswami G, Hartl C, et al. Shared molecular neuropathology across major psychiatric disorders parallels polygenic overlap. Science (New York, NY). 2018;359(6376):693.

93. Kurtz J-E, Ray-Coquard I. PI3 Kinase Inhibitors in the Clinic: An Update. Anticancer Research. 2012;32(7):2463–70.

94. Crabbe T. Exploring the potential of PI3K inhibitors for inflammation and cancer. Biochemical Society transactions. 2007;35(Pt 2):253–6.

95. Ito K, Caramori G, Adcock IM. Therapeutic potential of phosphatidylinositol 3-kinase inhibitors in inflammatory respiratory disease. The Journal of pharmacology and experimental therapeutics. 2007;321(1):1–8.

96. Kalkman HO. The role of the phosphatidylinositide 3-kinase-protein kinase B pathway in schizophrenia. Pharmacology & therapeutics. 2006;110(1):117–34.

97. McGuire JL, Hammond JH, Yates SD, Chen D, Haroutunian V, Meador-Woodruff JH, et al. Altered serine/threonine kinase activity in schizophrenia. Brain research. 2014;1568:42–54.

98. Enriquez-Barreto L, Morales M. The PI3K signaling pathway as a pharmacological target in Autism related disorders and Schizophrenia. Molecular and Cellular Therapies. 2016;4:2.

99. Kalkman HO. The role of the phosphatidylinositide 3-kinase–protein kinase B pathway in schizophrenia. Pharmacology & therapeutics. 2006;110(1):117–34.

100. Law AJ, Wang Y, Sei Y, O'Donnell P, Piantadosi P, Papaleo F, et al. Neuregulin 1-ErbB4-PI3K signaling in schizophrenia and phosphoinositide 3-kinase–p110delta inhibition as a potential therapeutic strategy. Proceedings of the National Academy of Sciences of the United States of America. 2012;109(30):12165–70.

101. Cheatham B, Vlahos CJ, Cheatham L, Wang L, Blenis J, Kahn CR. Phosphatidylinositol 3-kinase activation is required for insulin stimulation of pp70 S6 kinase, DNA synthesis, and glucose transporter translocation. Molecular and cellular biology. 1994;14(7):4902–11.

102. Dixon LB, Lehman AF, Levine J. Conventional antipsychotic medications for schizophrenia. Schizophrenia bulletin. 1995;21(4):567–77.

103. Tsygankov BD, Agasarain EG, Zykova AS. [Antipsychotic drugs and their influence on the carbohydrate metabolism in patients with schizophrenia-spectrum disorders]. Zhurnal nevrologii i psikhiatrii imeni SS Korsakova. 2014;114(5):86–91.

104. Davies JA. Mechanisms of action of antiepileptic drugs. Seizure. 1995;4(4):267–71.

105. Citrome L. Schizophrenia and valproate. Psychopharmacology bulletin. 2003;37 Suppl 2:74–88.

106. Schwarz C, Volz A, Li C, Leucht S. Valproate for schizophrenia. The Cochrane database of systematic reviews. 2008(3):Cd004028.

107. Rosenberg G. The mechanisms of action of valproate in neuropsychiatric disorders: can we see the forest for the trees? Cellular and Molecular Life Sciences. 2007;64(16):2090–103.

108. Juge-Aubry CE, Hammar E, Siegrist-Kaiser C, Pernin A, Takeshita A, Chin WW, et al. Regulation of the transcriptional activity of the peroxisome proliferator-activated receptor alpha by phosphorylation of a ligand-independent trans-activating domain. The Journal of biological chemistry. 1999;274(15):10505–10.

109. Wang L, Waltenberger B, Pferschy-Wenzig E-M, Blunder M, Liu X, Malainer C, et al. Natural product agonists of peroxisome proliferator-activated receptor gamma (PPARγ): a review. Biochemical Pharmacology. 2014;92(1):73–89.

110. Aljada A, Garg R, Ghanim H, Mohanty P, Hamouda W, Assian E, et al. Nuclear factor-kappaB suppressive and inhibitor-kappaB stimulatory effects of troglitazone in obese patients with type 2 diabetes: evidence of an antiinflammatory action? The Journal of clinical endocrinology and metabolism. 2001;86(7):3250–6.

111. Henry RR. Effects of troglitazone on insulin sensitivity. Diabetic medicine: a journal of the British Diabetic Association. 1996;13(9 Suppl 6):S148–50.

112. #xe9, rez M, #xed, Jos a, #xe9, Quintanilla RA. Therapeutic Actions of the Thiazolidinediones in Alzheimer&#x2019;s Disease. PPAR Research. 2015;2015:8.

113. Jiang L-Y, Tang S-S, Wang X-Y, Liu L-P, Long Y, Hu M, et al. PPARγ Agonist Pioglitazone Reverses Memory Impairment and Biochemical Changes in a Mouse Model of Type 2 Diabetes Mellitus. CNS Neuroscience & Therapeutics. 2012;18(8):659–66.

114. Heneka MT, Fink A, Doblhammer G. Effect of pioglitazone medication on the incidence of dementia. Annals of Neurology. 2015;78(2):284–94.

115. Masciopinto F, Di Pietro N, Corona C, Bomba M, Pipino C, Curcio M, et al. Effects of long-term treatment with pioglitazone on cognition and glucose metabolism of PS1-KI, 3xTg-AD, and wild-type mice. Cell Death Dis. 2012;3:e448.

116. Papadopoulos P, Rosa-Neto P, Rochford J, Hamel E. Pioglitazone Improves Reversal Learning and Exerts Mixed Cerebrovascular Effects in a Mouse Model of Alzheimer’s Disease with Combined Amyloid-β and Cerebrovascular Pathology. PloS one. 2013;8(7):e68612.

117. Sato T, Hanyu H, Hirao K, Kanetaka H, Sakurai H, Iwamoto T. Efficacy of PPAR-gamma agonist pioglitazone in mild Alzheimer disease. Neurobiology of aging. 2011;32(9):1626–33.

118. Sauerbeck A, Gao J, Readnower R, Liu M, Pauly JR, Bing G, et al. Pioglitazone attenuates mitochondrial dysfunction, cognitive impairment, cortical tissue loss, and inflammation following traumatic brain injury. Experimental Neurology. 2011;227(1):128–35.

119. Smith RC, Jin H, Li C, Bark N, Shekhar A, Dwivedi S, et al. Effects of pioglitazone on metabolic abnormalities, psychopathology, and cognitive function in schizophrenic patients treated with antipsychotic medication: a randomized double-blind study. Schizophrenia research. 2013;143(1):18–24.

120. Duncan GE, Moy SS, Lieberman JA, Koller BH. Typical and atypical antipsychotic drug effects on locomotor hyperactivity and deficits in sensorimotor gating in a genetic model of NMDA receptor hypofunction. Pharmacol Biochem Behav. 2006;85(3):481–91.

121. Duncan GE, Moy SS, Lieberman JA, Koller BH. Effects of haloperidol, clozapine, and quetiapine on sensorimotor gating in a genetic model of reduced NMDA receptor function. Psychopharmacology. 2006;184(2):190–200.

122. Fradley RL, O'Meara GF, Newman RJ, Andrieux A, Job D, Reynolds DS. STOP knockout and NMDA NR1 hypomorphic mice exhibit deficits in sensorimotor gating. Behav Brain Res. 2005;163(2):257–64.

123. Halene TB, Ehrlichman RS, Liang Y, Christian EP, Jonak GJ, Gur TL, et al. Assessment of NMDA receptor NR1 subunit hypofunction in mice as a model for schizophrenia. Genes Brain Behav. 2009;8(7):661–75.

124. Lee MA, Jayathilake K, Meltzer HY. A comparison of the effect of clozapine with typical neuroleptics on cognitive function in neuroleptic-responsive schizophrenia. Schizophrenia research. 1999;37(1):1–11.

125. Bhattacharya K. Cognitive Function in Schizophrenia: A Review. J Psychiatry Neurosci. 2015;18(187).

126. Baddeley A, Della Sala S, Papagno C, Spinnler H. Dual-task performance in dysexecutive and nondysexecutive patients with a frontal lesion. Neuropsychology. 1997;11(2):187–94.

127. Miyake A, Friedman NP, Emerson MJ, Witzki AH, Howerter A, Wager TD. The unity and diversity of executive functions and their contributions to complex "Frontal Lobe" tasks: a latent variable analysis. Cognitive psychology. 2000;41(1):49–100.

128. Rabin RA, Zakzanis KK, George TP. The effects of cannabis use on neurocognition in schizophrenia: a meta-analysis. Schizophrenia research. 2011;128(1-3):111–6.

129. Yücel M, Bora E, Lubman DI, Solowij N, Brewer WJ, Cotton SM, et al. The Impact of Cannabis Use on Cognitive Functioning in Patients With Schizophrenia: A Meta-analysis of Existing Findings and New Data in a First-Episode Sample. Schizophrenia bulletin. 2012;38(2):316–30.

130. Goldberg TE, Green MF. Neurocognitive functioning in patients with schizophrenia: an overview; in Neuropsychopharmacology - fifth generation of progress (eds) KL Davis, D Charney, JT Coyle2002. 657–69 p.

131. Kuleshov MV, Jones MR, Rouillard AD, Fernandez NF, Duan Q, Wang Z, et al. Enrichr: a comprehensive gene set enrichment analysis web server 2016 update. Nucleic acids research. 2016;44(W1):W90–7.

132. Iorio F, Rittman T, Ge H, Menden M, Saez-Rodriguez J. Transcriptional data: a new gateway to drug repositioning? Drug Discovery Today. 2013;18(7):350–7.

133. Roberts RC, Roche JK, Conley RR, Lahti AC. Dopaminergic synapses in the caudate of subjects with schizophrenia: relationship to treatment response. Synapse (New York, NY). 2009;63(6):520–30.

134. Chen EY, Tan CM, Kou Y, Duan Q, Wang Z, Meirelles GV, et al. Enrichr: interactive and collaborative HTML5 gene list enrichment analysis tool. BMC bioinformatics. 2013;14:128.

135. Milenkovic M, Mielnik CA, Ramsey AJ. NMDA receptor-deficient mice display sexual dimorphism in the onset and severity of behavioural abnormalities. Genes, brain, and behavior. 2014;13(8):850–62.

136. Moy SS, Nadler JJ, Young NB, Perez A, Holloway LP, Barbaro RP, et al. Mouse behavioral tasks relevant to autism: phenotypes of 10 inbred strains. Behav Brain Res. 2007;176(1):4–20.

137. Mielnik CA, Horsfall W, Ramsey AJ. Diazepam improves aspects of social behaviour and neuron activation in NMDA receptor-deficient mice. Genes, brain, and behavior. 2014;13(7):592–602.

